# Adjuvant Delivery Method and Nanoparticle Charge Influence Peptide Amphiphile Micelle Vaccine Bioactivity

**DOI:** 10.1101/2024.06.10.598369

**Authors:** Rui Zhang, Brett T. Rygelski, Luke E. Kruse, Josiah D. Smith, Xiaofei Wang, Brittany N. Allen, Jake S. Kramer, Gracen F. Seim, Trent J. Faulkner, Huihui Kuang, Efrosini Kokkoli, Adam G. Schrum, Bret D. Ulery

**Affiliations:** Department of Chemical and Biomedical Engineering, University of Missouri, Columbia, MO 65211; Department of Biochemistry, University of Missouri, Columbia, MO 65211; Institute for NanoBioTechnology, Johns Hopkins University, Baltimore, MD 21218; Department of Chemical and Biomolecular Engineering, Johns Hopkins University, Baltimore, MD 21218; Department of Molecular Microbiology and Immunology, University of Missouri, Columbia, MO 65211; Department of Surgery, University of Missouri, Columbia, MO 65211; NextGen Precision Health Institute, University of Missouri, Columbia, MO 65211; Materials Science and Engineering Institute, University of Missouri, Columbia, MO 65211

**Author notes:** To whom correspondence should be addressed: Bret Ulery, W2027 Lafferre Hall, 416 S. 6^th^ Street, Columbia, MO 65211. These authors contributed equally (Rui Zhang, Brett T. Rygelski).

**Keywords:** Subunit Vaccines, Peptide Amphiphile Micelles, Adjuvant, Co-Localization, CpG-B

## Abstract

Vaccines are an indispensable public health measure that have enabled the eradication, near elimination, and prevention of a variety of pathogens. As research continues and our understanding of immunization strategies develops, subunit vaccines have emerged as exciting alternatives to existing whole vaccine approaches. Unfortunately, subunit vaccines often possess weak antigenicity, requiring delivery devices and adjuvant supplementation to improve their utility. Peptide amphiphile micelles have recently been shown to function as both delivery devices and self-adjuvanting systems that can be readily associated with molecular adjuvants to further improve vaccine-mediated host immunity. While promising, many “design rules” associated with the plethora of underlying adjustable parameters in the generation of a peptide amphiphile micelle vaccine have yet to be uncovered. This work explores the impact micellar adjuvant complexation method and incorporated antigen type have on their ability to activate dendritic cells and induce antigen specific responses. Interestingly, electrostatic complexation of CpG to micelles resulted in improved *in vitro* dendritic cell activation over hydrophobic association and antigen|adjuvant co-localization influenced cell-mediated, but not antibody-mediated immune responses. These exciting results complement those previously published to build the framework of a micelle vaccine toolbox that can be leveraged for future disease-specific formulations.

## Introduction

Vaccines have become an important tool in modern medicine in the fight against a variety of infectious diseases. Recently, research has focused on using subunit vaccines, in part, due to their cost effectiveness and temperature stability making them cheaper and easier to employ in developing countries. Scientists hope to utilize subunit vaccines to fight a variety of diseases around the world but must overcome their inherent limitations including weak immunogenicity that has led to the need for delivery vehicles capable of improving the transportation of antigen and/or adjuvant to lymphoid organs of interest. One subunit vaccination approach is to utilize micelles as a vaccine carrier system (1–7). While micelles can be fabricated from a number of materials, peptide amphiphiles (PAs) are unique as they are comprised of hydrophilic peptide(s) conjugated to hydrophobic lipid(s) that can micellize in water (8–16) enabling this delivery technology to function as the subunit vaccine itself. Foundational research has shown that these self-assembling peptide amphiphile micelles (PAMs) are able to self-adjuvant incorporated antigens making them quite attractive synthetic vaccine candidates capable of inducing both cell-mediated and antibody-mediated immune responses (17,18). More recently, we have uncovered that transitioning from traditional diblock PAs to novel triblock PAs possessing a zwitterion-like peptide domain allows for the generation of novel PAM aggregates (Z-PAMs) which possess greatly altered vaccination potential (19,20).

PAM vaccines can be further enhanced through micelle-based adjuvant co-delivery (21). Specifically, molecular adjuvants hold promise as they have been widely utilized in subunit vaccine development against a number of emerging diseases including HIV, cancer, and influenza (22–30). One group of molecular adjuvants of particular interest are CpG oligodeoxynucleotides (CpG) which are unmethylated single stranded DNA that mimic bacterial genomic structure and stimulate toll like receptor 9 (TLR9) located in the endosomes of antigen presenting cells (31,32). CpG ODN 1826 is a particularly promising adjuvant as it can enhance both cell-mediated and antibody-mediated immune responses (33–37). The multipronged activation potential of CpG allows for it to be utilized to convey host protection against intracellular and extracellular pathogens (38–45). Unfortunately, like many other nucleic acid-based products, CpG is a large (∼ 6.4 kDa), negatively charged molecule limiting its capacity to be readily internalized and processed by antigen presenting cells (APCs). Moreover, as interstitially administrated soluble CpG preferentially disseminates into blood vessels and accumulates in the liver and spleen (46), cell-mediated immune responses (*i.e.*, cytotoxic T cell and helper T cell engagement) are often hindered. Thus, an effective delivery vehicle is often required to help CpG reach its desirable site of action and initiate productive immune responses (47–61).

Previous PAM antigen|adjuvant co-localization has been achieved by leveraging the hydrophobic micelle core to directly associate lipid-based molecular adjuvants (*i.e.*, MPLA and Pam_2_C) (21,62). While promising, some adjuvants like CpG do not possess the hydrophobic domain necessary to facilitate this means of co-localization. Therefore, either the adjuvant needs to be modified to facilitate hydrophobic association or the third PA block altered to facilitate charge complexation on the PAM surface. Expanding on previous PAM vaccine research (17–21,62,63), this paper explores the efficacy ofdifferent methods of CpG adjuvant co-localization with PAM-associated subunit vaccines, specifically focusing on a cytotoxic T cell antigen and a linked-recognition antigen possessing both helper T cell and B cell epitopes. These efforts coupled with earlier results further develop design rules for use in disease-specific vaccines that will be developed in the future.

## Materials and Methods

### Peptide, Peptide Amphiphile, and Lipidated CpG Synthesis and Purification

A model cytotoxic T cell antigen containing peptide (*i.e.*, EQLESIINFEKLTE - OVA_CytoT_) was modified to create K-OVA_CytoT_-K_8_ (cationic peptide) and K-OVA_CytoT_-(KE)_4_ (zwitterion-like peptide) employing Rink Amide on-resin synthesis with Fmoc protection on the terminal lysine side chains using a Tetras Peptide Synthesizer housed in the Molecular Interactions Core at the University of Missouri. The same approach was taken for the synthesis of OVA_BT_ (ESLKKISQAVHAAHAEINEAGRE) yielding K-OVA_BT_-K_8_ and K-OVA_BT_-(KE)_4_. The Fmoc protection groups were removed using 25% piperidine in dimethylformamide (DMF) incubated for 15 minutes at room temperature (RT) while being shaken. Two palmitic acids were then added to the peptides to yield four peptide amphiphiles (PAs) - P_2_K-OVA_CytoT_-K_8_ (OVA_CytoT_ Cat-PA), P_2_K-OVA_CytoT_-(KE)_4_ (OVA_CytoT_ Z-PA), P_2_K-OVA_BT_-K_8_ (OVA_BT_ Cat-PA), and P_2_K-OVA_BT_-(KE)_4_ (OVA_BT_ Z-PA). These reactions were carried out in N-methyl-2-pyrrolidone (NMP) with the following component ratios: 10x molar excess palmitic acid, 11x molar excess hydroxybenzotriazole (HOBt), 9.9x molar excess 2-(1h-benzotriazole-1-yl)-1,1,3,3-tetramethyluronium hexafluorophosphate (HBTU), and 10x molar excess N,N-diisopropylethylamine (DIEA) (all procured from Sigma-Aldrich) in comparison to the quantity of peptide. Palmitic acid attachment for PA synthesis was confirmed via the Kaiser test where ninhydrin shows the presence of deprotected primary amines by turning the solution blue. After palmitic acid addition, PAs were cleaved from resin by incubating at RT for 2 hours in the following solution: 90% trifluoracetic acid (TFA), 2.5% triisopropylsilane (TIS), 2.5% deionized, distilled water (ddH_2_O), 2.5% ethanedithiol (EDT), and 2.5% thioanisole (TA) supplemented with 3.3% w/w phenol (all purchased from Sigma-Aldrich). After cleavage, 20 mL of diethyl ether was added to the solution to facilitate PA precipitation followed by centrifugation at 2,500 x g for 3 minutes and decantation with the pellet being saved for further use. This centrifugation step was repeated for a total of 3 times after which the PAs were lyophilized in 50% water and 50% acetonitrile. After freeze-drying, the PAs were purified and analyzed using mass-spectrometry controlled high pressure liquid chromatography (LCMS, Beckman System Coulter Gold) in the Molecular Interactions Core at the University of Missouri, after which PAs were lyophilized into 1 mg aliquots. OVA_CytoT_ and OVA_BT_ peptide were also synthesized and purified using the same process but with the N-terminus of the peptides acetylated using a NMP solution containing 5% acetic anhydride and 7% DIEA. Lipidated CpG molecules were synthesized by covalently conjugating CpG ODN 1826 (purchased from Trilink Biotechnologies) to a dialkyl C16 (diC_16_) tail. The diC_16_ hydrophobic tail was synthesized by first reacting hexadecanol to L-glutamic acid (linker between the two C_16_ chains), followed by the addition of succinic anhydride that acts as a short spacer between the tail and CpG headgroup (64,65). The conjugation, purification and characterization of the CpG amphiphiles was done as reported previously (66–68). Briefly, CpG with an amine-terminated, six-carbon spacer on either the 5’ or 3’ end (*i.e.*, NH_2_-C6-CpG or CpG-C6-NH_2_) was conjugated to the diC_16_ tail through amide coupling to yield diC_16_-CpG (Lipid-CpG 5’) or CpG-diC_16_ (Lipid-CpG 3’), respectively. The unreacted CpG molecules were separated from the CpG amphiphiles using reverse phase high performance liquid chromatography (RP-HPLC). The molecular weights of the CpG amphiphiles were verified via liquid chromatography-mass spectroscopy (LC-MS). The concentration of the amphiphiles was measured by absorbance at 260 nm.

### Transmission Electron Microscopy

Micelle formation and architecture were confirmed using transmission electron microscopy (TEM). PA solutions in ddH_2_O (10 μM) were generated in microcentrifuge tubes. Additional samples adding 0.25 μM CpG ODN 1826 (CpG, Trilink Biotechnology, San Diego, CA) and 0.25 μM lipidated CpG to the PAs were utilized to determine if their association with the micellar structure affected nanoparticle size and shape. The addition of CpG allows for the electrostatic complexation of the negatively charged CpG DNA to the micelles while the addition of lipidated CpG allows for hydrophobic association of the CpG (**Scheme 1**). After the samples were prepared, carbon grids were first glow discharged using a Pelco Easiglow, after which 5 μL of 10 μM PA solutions were added to the TEM grids, allowed to equilibrate for 5 minutes, and then the liquid was wicked away. Nanotungsten stain (5 μL) was added to the grids and allowed to incubate for 5 minutes. After the nanotungsten stain was wicked away, the grids were assessed using a JEOL JEM-1400 (housed in the University of Missouri’s Electron Microscopy Core).

**Scheme 1.**
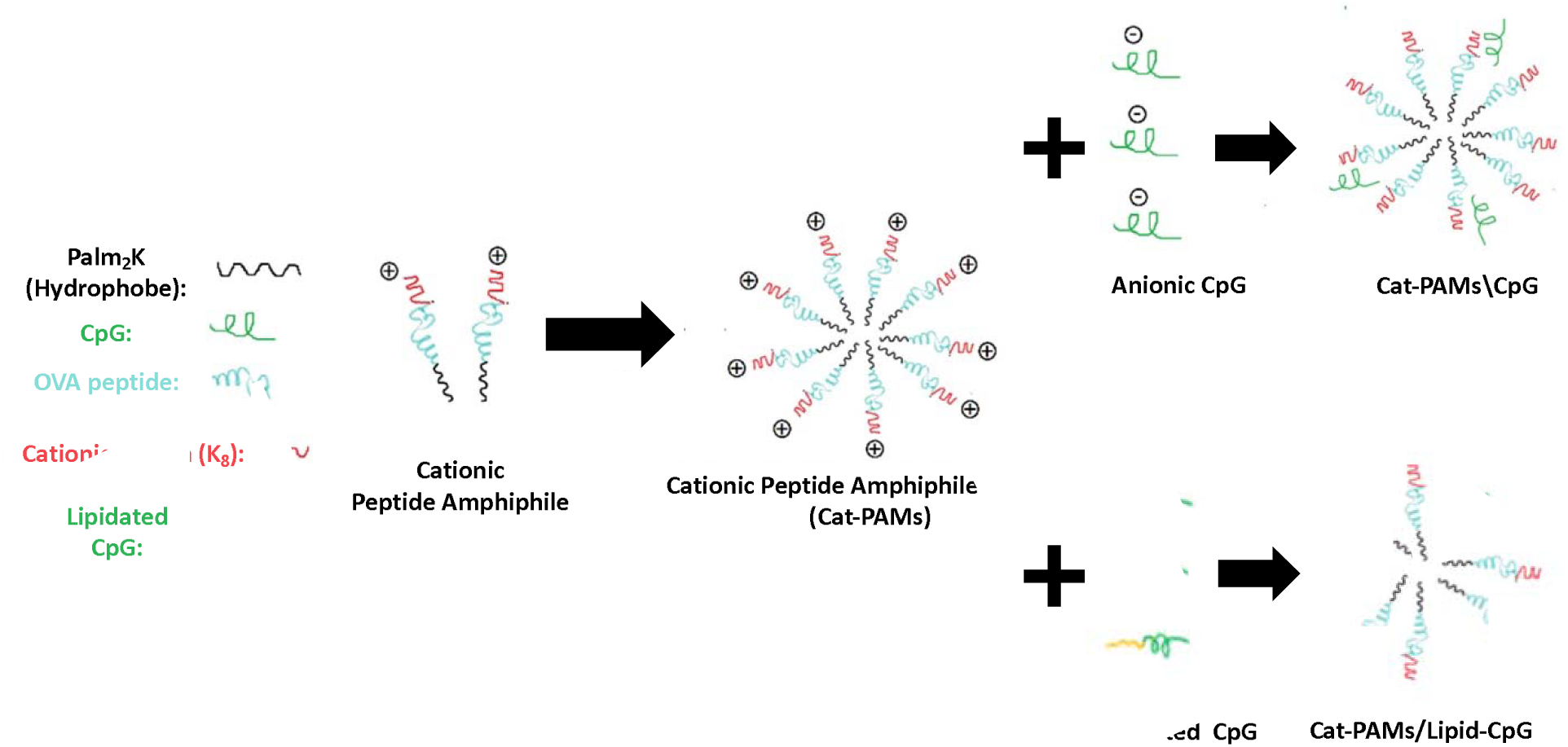
Fabrication of electrostatically complexed and hydrophobically associated Cat-PAM\CpG and Cat-PAM/Lipid-CpG, respectively. A cationic region (*i.e.*, K_8_) is included during the PA synthesis process which yields cationic PAs. After self-assembly, Cat-PAMs possess positive surface charge that enables them to electrostatically complex anionic CpG to their corona. Alternatively, lipidation of CpG on the 3’ or 5’ end allows for its association with the hydrophobic moiety of the peptide amphiphile during micellization.

### Preparation and Activation of Bone Marrow-Derived Dendritic Cells

C57BL/6J and Balb/c mouse femurs and tibias were harvested, from which cells were collected by flushing the bone marrow with complete RPMI media (RPMI 1640 media - Sigma-Aldrich) supplemented with 10% fetal bovine serum - FBS, 1% penicillin-streptomycin, and 50 μM β-mercaptoethanol) that was passed through a cell strainer (70 μm mesh size). Red blood cells were lysed by ammonium-chloride-potassium (ACK) lysing buffer before stromal cells were seeded on non-tissue culture treated petri-dishes. These stromal cells were then cultured in bone marrow dendritic cell (BMDC) differentiation media (complete RPMI supplemented with 20 ng/mL granulocyte-macrophage colony-stimulating factor (GM-CSF)) at 37 °C with 5% CO_2_ for which culture media was changed on days 3, 6, and 8. Stromal cells were considered to have differentiated into BMDCs after 10 days of incubation and were seeded in 24-well plates at 10^5^ cells/well, and allowed to incubate overnight. The cells were exposed to 100 μL of phosphate buffered saline (PBS) alone or with one of a variety of different controls, immunostimulants, and vaccines including peptide, PAMs, CpG, lipidated CpG, or combinations of these molecules (Tables I - II). This was supplemented with 400 μL of complete RPMI media and the cells were then allowed to incubate for 24 hours. Treated BMDCs were harvested, blocked with anti-CD16/32 for 10 minutes, and stained with fluorescently-labeled antibodies (*i.e.,* PE/Cy7-CD11c, APC-MHC II, PE-CD86, and FITC-CD40 – Biolegend) for 30 minutes. Stained cells were then fixed with 4% paraformaldehyde and analyzed by flow cytometry.

**Table I.** Therapeutic formulations for the *in vitro* OVA_CytoT_ BMDC activation study.

| BMDC Stimulus Groups | Dose |
| --- | --- |
| PBS | Phosphate Buffered Saline |
| OVA<sub>CytoT</sub> Peptide | 180 pmol OVA<sub>CytoT</sub> Peptide |
| OVA<sub>CytoT</sub> Cat-PAMs | 180 pmol Palm<sub>2</sub>K-OVA<sub>CytoT</sub>-K<sub>8</sub> |
| OVA<sub>CytoT</sub> Z-PAMs | 180 pmol Palm<sub>2</sub>K-OVA<sub>CytoT</sub>-(KE)<sub>4</sub> |
| CpG | 4.5 pmol CpG |
| Lipid-CpG 5' | 4.5 pmol Lipid-CONH-C6-CpG |
| Lipid-CpG 3' | 4.5 pmol CpG-C6-NHCO-Lipid |
| OVA<sub>CytoT</sub> peptide + CpG | 180 pmol OVA<sub>CytoT</sub> peptide<br>4.5 pmol CpG |
| OVA<sub>CytoT</sub> peptide + Lipid-CpG 5' | 180 pmol OVA<sub>CytoT</sub> peptide<br>4.5 pmol Lipid-CONH-C6-CpG |
| OVA<sub>CytoT</sub> peptide + Lipid-CpG 3' | 180 pmol OVA<sub>CytoT</sub> peptide<br>4.5 pmol CpG-C6-NHCO-Lipid |
| OVA<sub>CytoT</sub> Cat-PAMs\CpG | 180 pmol Palm<sub>2</sub>K-OVA<sub>CytoT</sub>-K<sub>8</sub><br>4.5 pmol CpG |
| OVA<sub>CytoT</sub> Cat-PAMs/Lipid-CpG 5' | 180 pmol Palm<sub>2</sub>K-OVA<sub>CytoT</sub>-K<sub>8</sub><br>4.5 pmol Lipid-CONH-C6-CpG |
| OVA<sub>CytoT</sub> Cat-PAMs/Lipid-CpG 3' | 180 pmol Palm<sub>2</sub>K-OVA<sub>CytoT</sub>-K<sub>8</sub><br>4.5 pmol CpG-C6-NHCO-Lipid |
| OVA<sub>CytoT</sub> Z-PAMs\CpG | 180 pmol Palm<sub>2</sub>K-OVA<sub>CytoT</sub>-(KE)<sub>4</sub><br>4.5 pmol CpG |
| OVA<sub>CytoT</sub> Z-PAMs/Lipid-CpG 5' | 180 pmol Palm<sub>2</sub>K-OVA<sub>CytoT</sub>-(KE)<sub>4</sub><br>4.5 pmol Lipid-CONH-C6-CpG |
| OVA<sub>CytoT</sub> Z-PAMs/Lipid-CpG 3' | 180 pmol Palm<sub>2</sub>K-OVA<sub>CytoT</sub>-(KE)<sub>4</sub><br>4.5 pmol CpG-C6-NHCO-Lipid |

**Table II.**
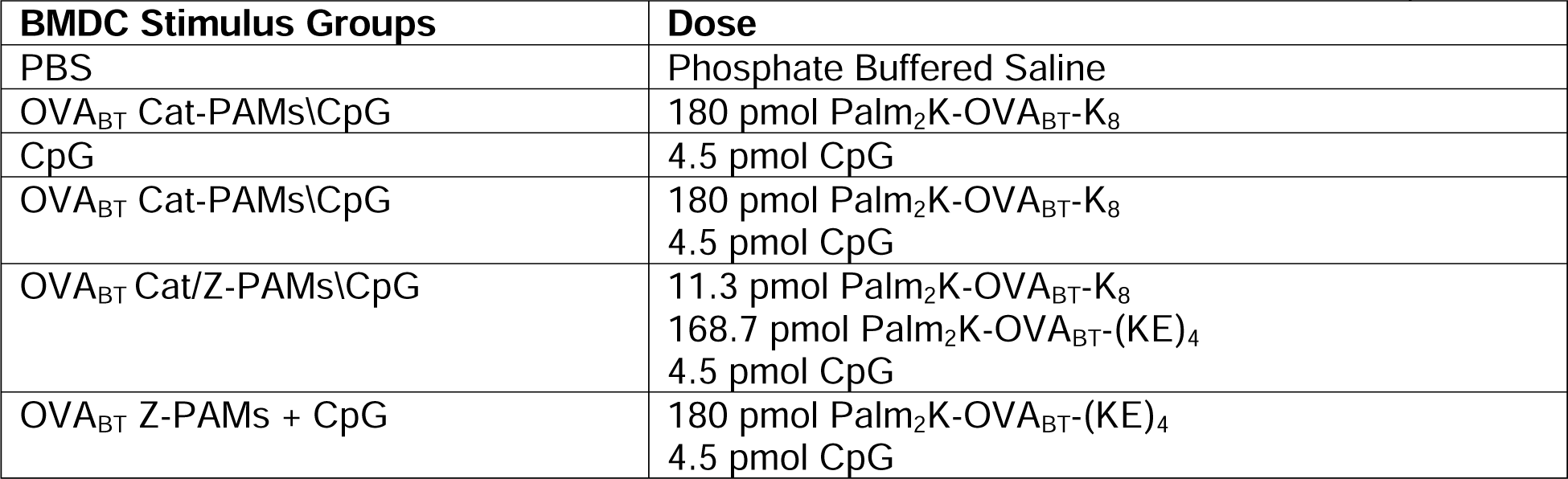
Therapeutic formulations for the *in vitro* OVA_BT_ BMDC activation study.

### Förster Resonance Energy Transfer

Förster Resonance Energy Transfer (FRET) was used to confirm co-localization of PAMs and CpG or the lack thereof. The FRET pair utilized for this study included FAM (5,6-carboxyfluoroscein) and TAMRA (5-carboxytetramethylrhodamine). FAM modified CpG (*i.e.*, FAM-CpG) was purchased from Trilink Bio-Technology (San Diego, CA). PAs (19.92 μL of a 301 μM solution) were mixed with TAMRA (0.3 μL of a 100 μM solution) fluorophore in a 300 μL methanol (MeOH) solution and film dried utilizing nitrogen gas. The dried PA/TAMRA film was then hydrated in ddH_2_O to create 40 μM solutions with PAM core entrapped hydrophobic TAMRA. FAM-CpG was mixed with non-fluorescent CpG at a 1:70 molar ratio of which 0.267 pmol in 3.375 μL was added to the TAMRA-loaded PA methanol solution before micelles were fabricated. Controls for this experiment included TAMRA-loaded PAMs and FAM-CpG alone. Fluorescence intensity was then measured for each formulation at RT using a BioTek Cytation 5 plate reader employing 2 nm intervals from 475 nm to 725 nm.

### Zeta Potential

PAMs (750 μL of a 10 μM ddH_2_O solution) were added to a new dynamic light scattering (DLS) polycarbonate cuvette with CpG added to the PAMs at a 1:40 molar ratio when applicable. A Zetasizer NanoZ system (Malvern Panalytical, Malvern, United Kingdom) was utilized to determine Zeta Potential employing the Hückel Method for proteins with no equilibration time. Zeta potential experiments were conducted in triplicates with each experiment consisting of five runs and three measurements taken per run.

### Vaccine-Induced Cell-Mediated Immunity

All *in vivo* vaccination studies were conducted using protocols approved by the University of Missouri Animal Care and Use Committee (ACUC). C57BL/6J mice (6 - 8 weeks old) were obtained from Jackson Laboratory with each vaccine group containing a group size consisting of at least 8 animals with a minimum of 4 males and 4 females each. Animals were vaccinated with the appropriate vaccine formulation (**Table III**) on Day 0 and Day 10 using 100 μL inoculations injected subcutaneously into the nape of the neck. On Day 20, the mice were sacrificed by carbon dioxide asphyxiation followed by cervical dislocation, after which the draining lymph nodes and spleen were harvested and kept on ice in 500 μL complete RPMI media.

**Table III.**
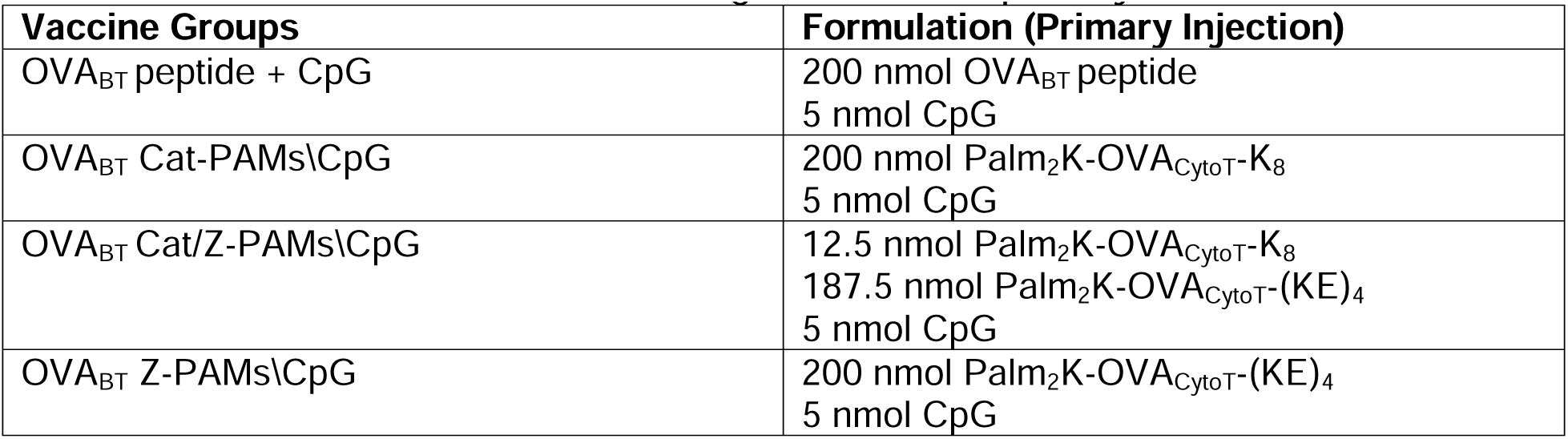
Vaccine formulations for the *in vivo* OVA_CytoT_ PAM immunogenicity study.

### Spleen Processing and Analysis

In a sterile biosafety cabinet, spleens were gently mashed with a tissue grinder in a 1.5 mL microcentrifuge tube, after which homogenate containing media was added to a 15 mL conical tube. After 5 mL of sterile PBS was added to the tube, the cells were centrifuged at 350 x g for 9 minutes at RT. The liquid was decanted, and the pellet dispersed by racking the bottom of the tube. PBS (10 mL) was added to the splenic single cell suspensions of which 200 μL was taken for tetramer staining to identify the percentage of cytotoxic T cells expressing a SIINFEKL-specific T cell receptor (*see Tetramer Staining section*). Cells were then centrifuged at 350 x g for 9 minutes at RT to form a pellet after which liquid decantation and pellet dispersion by racking were performed again. ACK lysing buffer (1 mL) was added to the cell suspension to lyse red blood cells and allowed to incubate for 3 minutes at RT. PBS (10 mL) was added to dilute the lysing buffer after which the cells were centrifuged at 350 x g for 9 minutes at RT. If the pellet appeared red, then the ACK lysing step was repeated to make sure all red blood cells were removed. Racking was used to disperse the pellet which was followed by the addition of 5 mL of complete RPMI media and filtration through a 70 μm cell strainer into a 50 mL conical tube. An additional 5 mL of complete RPMI media was added to the first conical tube and also passed through the filter to ensure high fidelity cell collection. Splenocytes were then counted using a Z2 Coulter Particle Count and Size Analyzer (Beckman Coulter, Brea, CA) to determine the total number of cells in each tube. Complete RPMI media was added to each tube to reach a desired concentration of 2 million cells per mL. Splenic cell solutions (500 μL containing 1 million cells) and antigen stimulus solutions (500 μL of media with 4 μg/mL OVA_CytoT_ peptide) were then added to 24 well plates. Samples for cytokine analysis (*see IFN-γ Staining section*) were exposed to Brefeldin A after 2 hours and incubated for a total of 6 hours.

### Lymph Node Processing and Analysis

The draining lymph nodes were processed similarly to spleens (*see Spleen Processing and Analysis section*) except without the ACK lysing buffer step as it is unnecessary due to the lack of red blood cells in this tissue. Lymphocytes were similarly counted, and the solution volume brought to a concentration of 2 million cells per mL using complete RPMI media from which 100,000 cells from each sample were taken for SIINFEKL-specific T cell receptor assessment (*see Tetramer Staining section*). Lymph node cell solutions (500 μL containing 1 million cells) were added to each well in 24 well plates and stimulated with antigen stimulus solutions (500 μL of media with 4 μg/mL OVA_CytoT_ peptide). Samples were incubated for cytokine analysis (*see IFN-γ Staining section*) according to the protocol outlined for splenic cells (*see Spleen Processing and Analysis section*) with Brefeldin added at the 2-hour mark and samples processed after 6 total hours.

### Tetramer Staining

Both splenocytes and lymphocytes were stained using allophycocyanin (APC) labeled MHC I haplotype matched SIINFEKL tetramer received from the NIH tetramer facility. After cells were transferred to flow cytometry tubes, 2 mL of PBS was added to each tube and kept on ice in order to maintain a temperature of 0 - 4 °C. Tubes were then centrifuged at 350 x g for 9 minutes at 4 °C after which they were decanted and returned to the ice bath. FACS buffer (3 mL) was added to each tube followed by centrifugation at 350 x g for 9 minutes at 4 °C. After decanting the FACS buffer, 100 μL of blocking buffer (1:100 dilution of TruStain anti-mouse CD16/32) was added to each tube and incubated for 5 minutes on ice. After blocking, 100 μL of staining solution containing APC SIINFEKL tetramer (1:600 dilution) and fluorescein isothiocyanate (FITC) CD8 antibody (1:100 dilution, Biolegend) was added to each tube in the ice bath and incubated for 30 minutes in the dark. FACS buffer (3 mL) was then added to each tube and centrifuged at 350 x g for 9 minutes at 4 °C. After decanting the solution, 200 μL of 4% paraformaldehyde fixation buffer was added to the tubes and incubated for 30 minutes at RT. FACS buffer (3 mL) was then added to each tube after which the cells were centrifuged at 350 x g for 9 minutes at 20 °C. The solution was decanted and then 3 mL of PBS was added to each tube. The tubes were centrifuged again at 350 x g for 9 minutes at 20 °C after which the solution was decanted once more. After adding 200 μL of PBS to each tube, the cells were analyzed via flow cytometry.

### IFN-γ Staining

For both splenocytes and lymphocytes, the medium from the well plates was removed and 0.5 mL of cell dissociation buffer (Gibco) was added to each well and allowed to incubate for 3 minutes at RT. The cell dissociation buffer was discarded and the plates were hit against their side to dislodge the cells that were then removed using multiple rounds of pipetting with 0.5 mL of PBS. Cells were added to flow tubes and processed similarly to the method previously described (*see Tetramer Staining section*). The major difference in approach is that after fixation, intracellular staining was performed. Permeabilization buffer (3 mL) was added to each tube and incubated for 10 minutes at RT before being centrifuged at 350 x g for 9 minutes at 20 °C. After the solution was decanted, 100 μL of staining solution containing 0.5 μg of APC IFN-γ antibody (Biolegend) in permeabilization buffer was added to each tube and incubated for 30 minutes in the dark at RT. An additional 2 mL of permeabilization buffer was added to each tube followed by centrifugation at 350 x g for 9 minutes at 20 °C. The solution was decanted and 2 mL of PBS was added to each tube followed by an additional centrifugation at 350 x g for 9 minutes at 20 °C. After the solution was decanted, 200 μL of PBS was added to each tube in preparation for running flow cytometry.

### Cell Uptake Assessment

RAW 264.7 (macrophage-like cells – MØs, ATCC) or naïve T cells isolated from lymph nodes (MojoSort Mouse CD4 Naïve T Cell isolation kit) were seeded at a density of 2 x 10^5^ cells/well on coverslips in a 24-well plate and incubated overnight. Cells were then exposed to different drug formulations with fluorescently labelled CpG (**Table IV**) for 1 hour at 37 °C with 5% CO_2_ followed by washing with PBS. Cells were then fixed with 4% paraformaldehyde and analyzed by flow cytometry.

**Table IV.**
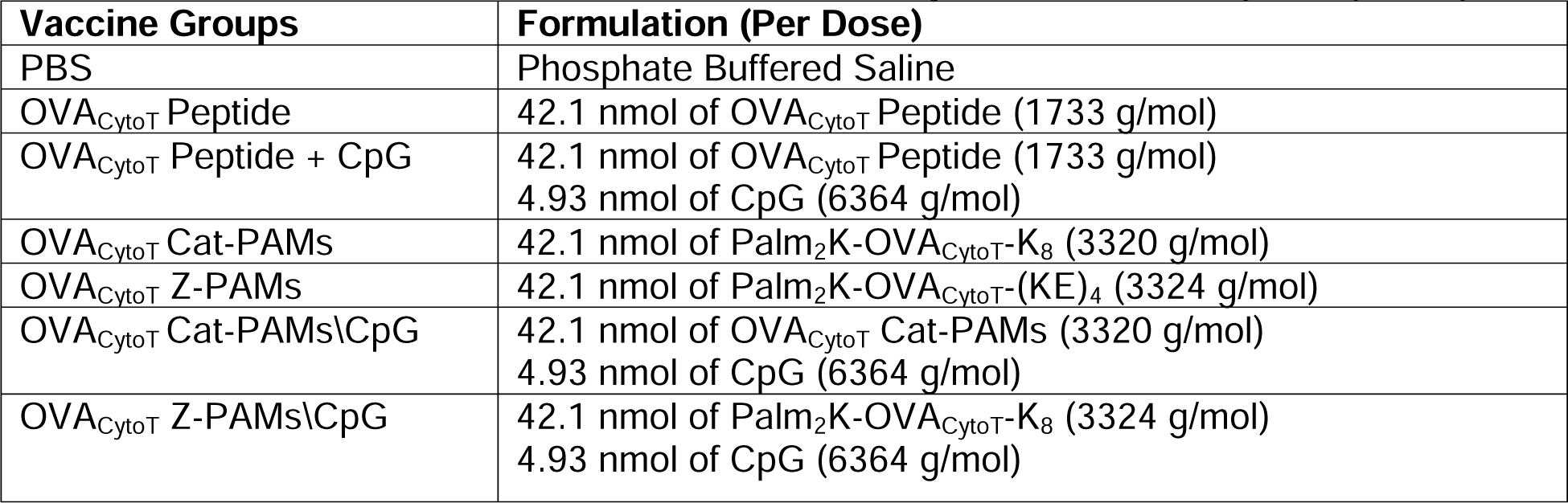
Therapeutic formulations for the *in vitro* OVA_BT_ BMDC cell uptake study.

### Flow Cytometry

All flow cytometry tubes were analyzed using a BD LSRFortessa X-20 flow cytometer or a Beckman Coulter CyAn ADP, both located in the Cell and Immunobiology Core at the University of Missouri. Cell counts were set to be greater than 10,000 for lymphocytes and more than 50,000 for splenocytes. Flow cytometry data was analyzed using Flow Jo software, first utilizing forward scatter versus side scatter analysis to generally separate out cell population. Further analysis allowed for the identification of CD11c^+^, MHC II^+^, CD86^+^, CD40^+^, CD8^+^, SIINFEKL^+^, and IFN-γ^+^ cell populations.

### Vaccine-Induced Antibody-Mediated Immunity

Sex-matched Balb/C mice (at least 4 males and 4 females each per group), 6 - 8 weeks old, were obtained from Jackson Laboratories and subcutaneously administrated different vaccine formulations (**Table V**) in the nape of the neck. Primary and boost inoculations were given at Day 0 and Day 28, respectively. Mice were sacrificed 14 days post booster vaccination (*i.e.*, Day 42) with their whole blood collected via cardiac puncture and centrifuged at 10,000 x g for 10 min to separate out the red blood cells. The resulting serum supernatants were harvested and stored at −80 °C until further analyzed. This *in vivo* experiment was performed according to a protocol approved by the Animal Care and Use Committee (ACUC) at the University of Missouri.

**Table V.**
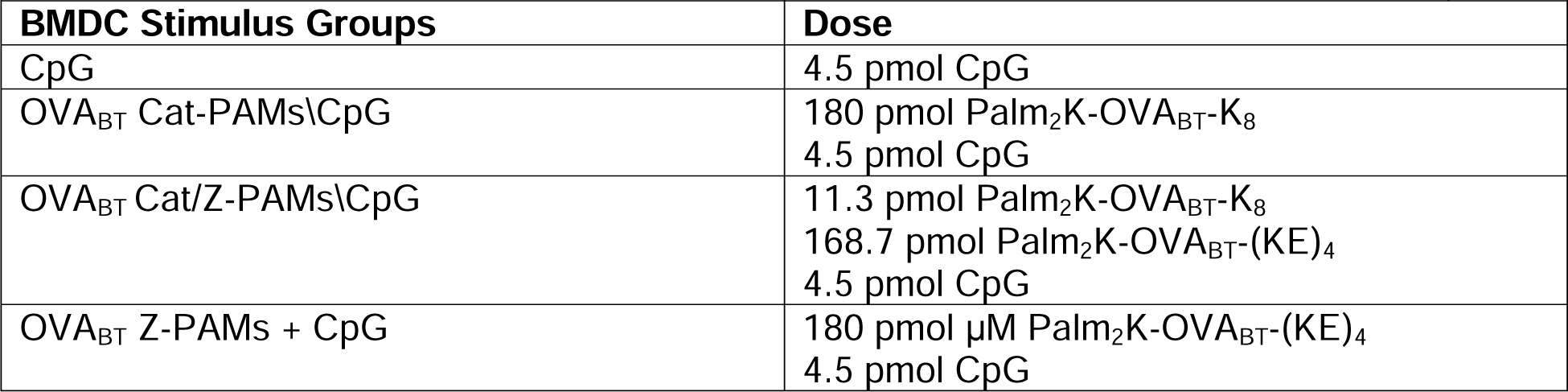
Therapeutic formulations for the *in vivo* OVA_BT_ PAM immunogenicity study. Booster vaccinations were half the dosage of the listed primary vaccination.

### Antibody Response Characterization

High binding, 96-well enzyme-linked immunosorbent assay (ELISA) plates (Santa Cruz Biotechnology) were coated overnight with 4 μg/mL OVA_BT_ peptide in PBS. Wells were washed with PBS-T (0.05% Tween-20 in PBS) and blocked with 10% FBS in PBS (blocking buffer) for 1 hour. Serum was serially diluted two-fold in blocking buffer across the plate and allowed to incubate for 2 hours. Wells were then washed with PBS-T and incubated with 1:3000 diluted detection antibody (*i.e.*, goat anti-mouse IgG(H&L), IgG1, IgG2a, or IgG3 – Thermo-Fisher) for 1 hour. After additional washing with PBS-T, wells were incubated for 30 mins with 100 μL TMB substrate (Biolegend) and optical density (OD) was defined as absorbance measured at 650 nm using a Biotek Cytation 5 spectrofluorometer. End-point antibody titers were defined as the greatest serum dilution where ELISA OD was at least twice that of serum from mice vaccinated with PBS alone. If end-point titers were not reached with one plate, then additional titrations were utilized until ODs were diluted below the cut-off limit.

### Lymphocyte Isolation, Antigenic Challenge, and Stimulus Assessment

Draining lymph nodes and spleens were harvested, processed, and investigated similarly to the protocols mentioned above. One major difference was that the restimulation experiment was carried out for 72 hours instead of 6 hours to allow for significant generation and secretion of cytokines. At this time point, supernatants were drawn off the cells and analyzed for cytokine content (*i.e.*, IL-2, IFN-γ, and TNF-α) by ELISA kits (Biolegend).

### Statistical Analysis

JMP pro software was used to compare groups employing an analysis of variance (ANOVA) assessment followed by Tukey’s honest significant differences (HSD) test in order to determine pairwise statistically significant differences between groups (p < 0.05). Within graphs, groups that possess different letters have statistically significant differences in mean whereas those that possess the same letter have statistically insignificant differences. Error bars indicate a single standard deviation from the mean.

## Results

### OVA_CytoT_ PA Micellization

In order to study whether PAMs can induce a cell-mediated immune response, a mildly immunogenic ovalbumin derived peptide, OVA_CytoT­_ (sequence EQLESIINFEKLTE), was employed as it is a known model cytotoxic T Cell antigen. OVA_CytoT­_ peptide was transformed into PAs through lipidation and non-native peptide block addition yielding OVA_CytoT_ Cat-PA (P_2_K-OVA_CytoT_-K_8_) and OVA_CytoT_ Z-PA (P_2_K-OVA_CytoT_-(KE)_4_). As changes in peptide sequence and charge can greatly influence material properties including micelle shape and size, experiments assessing these physical attributes were performed. Transmission electron microscopy (TEM) confirmed Cat-PAs and Z-PAs formed spherical and short cylindrical micelles (PAMs), respectively (**Figure 1**).

**Figure 1.**
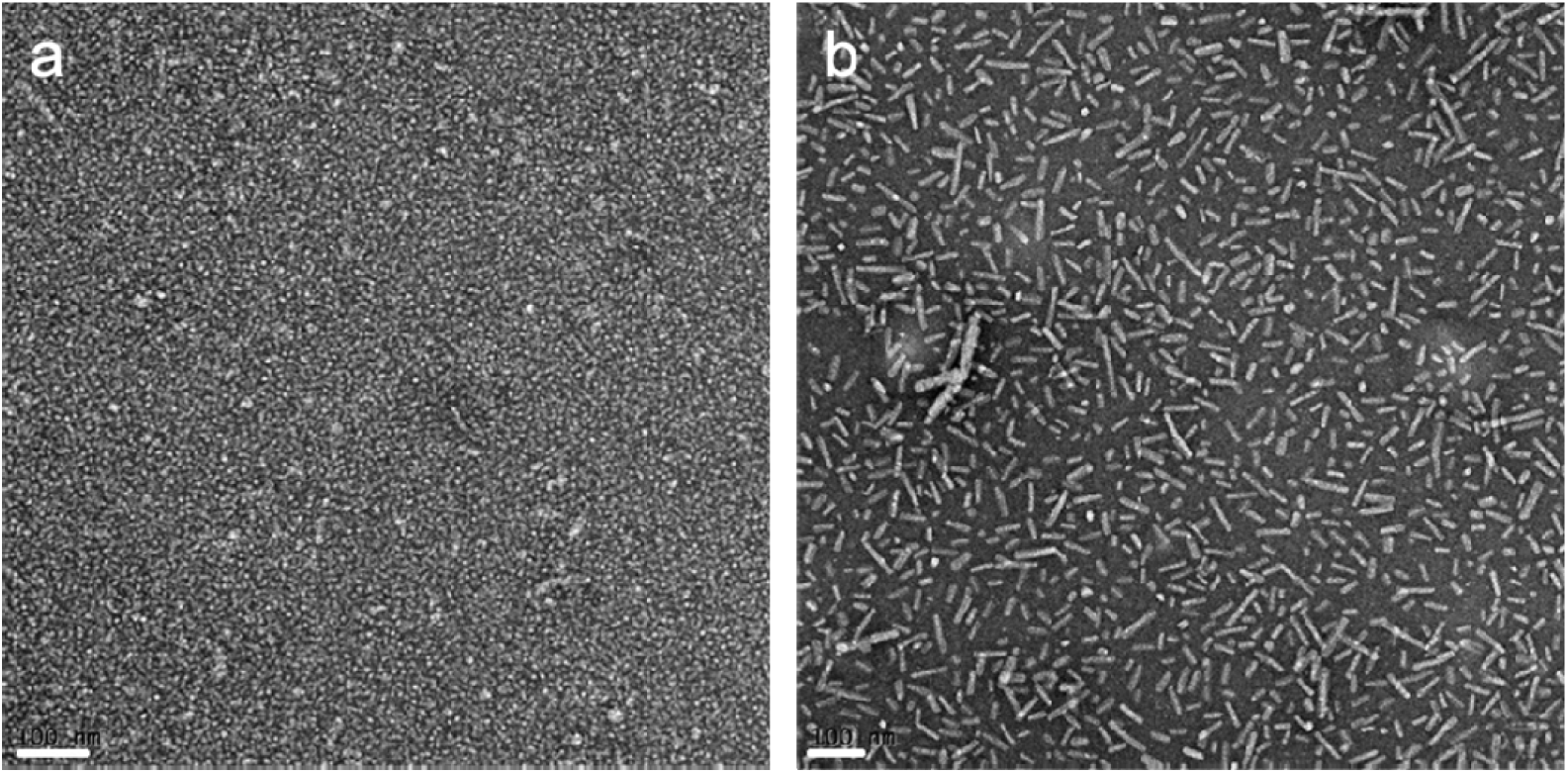
OVA_CytoT_ PAs readily form small micelles. Transmission electron microscopy (TEM) analysis shows the formation of **a** spherical micelles for OVA_CytoT_ Cat-PAMs and **b** short cylindrical micelles for OVA_CytoT_ Z-PAMs at 10 μM (scale bar = 100 nm). Representative images are shown.

### CpG Adjuvanticity

To determine the capacity of CpG to function as a localized adjuvant for PAM vaccination, a method for incorporating CpG with the PAM must be established. Two such colocalization strategies, electrostatic complexation and hydrophobic association, were assessed for their relative capacity to activate BMDCs *in vitro* as assessed by cell surface marker expression. Electrostatic complexation was hypothesized to occur between cationic PAMs and anionic CpG and the importance of the surface charge was assessed by employing a zwitterion-like PAM counterpart. To facilitate hydrophobic incorporation, CpG was lipidated at either the 5’ or 3’ terminus and added to PA solutions during the micelle fabrication process. Having verified that the presence of CpG did not alter micelle size and shape (**Figure S1**), each incorporation modality was evaluated for its capacity to alter adjuvant bioactivity. Vaccine formulations and their appropriate controls were incubated with BMDCs for 24 hours, after which activation-associated cell specific markers (*i.e.*, CD86 and MHC II) were stained and quantified by flow cytometry (**Figure 2**).

**Figure 2.**
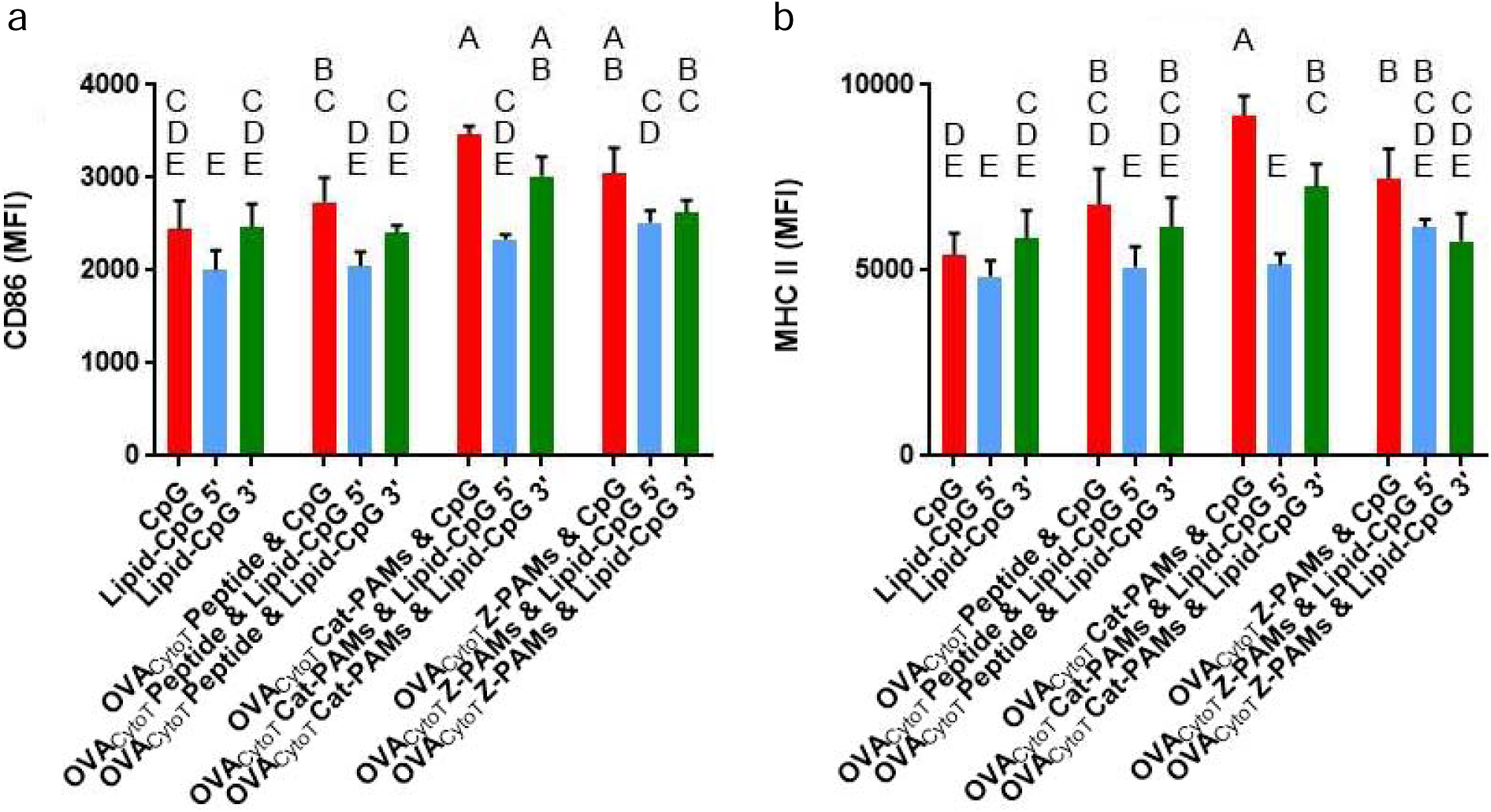
Lipidation and micelle incorporation method impact CpG-mediated BMDC activation. BMDCs were stimulated with the indicated formulations for 24 hours, after which cell activation was assessed by determining the quantity of cell surface **a** CD86 and **b** MHC II present. CpG (red), Lipid-CpG 5’ (blue) and Lipid-CpG 3’ (green) were compared alone, with peptide, and with PAMs. Within a graph, groups that possess different letters have statistically significant differences in mean (*p* ≤ 0.05) whereas those that possess the same letter do not.

No statistically significant differences in either CD86 or MHC II MFI were observed when peptide was co-delivered with unmodified or lipidated CpG when compared to their respective CpG alone, peptide free controls. In fact, the only distinguishable groups among these formulations were cells exposed to unmodified CpG performed better than those receiving 5’-lipidated CpG. Cells treated with OVA_CytoT_ Cat-PAMs & CpG were found to possess a significant enhancement in both CD86 and MHC II surface expression when compared to the CpG alone control and peptide plus CpG groups. Increased CD86 expression was observed over 5’-lipidated CpG associated with cationic PAMs (*i.e.*, OVA_CytoT_ Cat-PAMs & Lipid-CpG 5’) and both 5’- and 3’-lipidated CpG variants associated with zwitterionic PAMs (*i.e.*, OVA_CytoT_ Z-PAMs & Lipid-CpG 3’ and OVA_CytoT_ Z-PAMs & Lipid-CpG 5’). Additionally, cells exposed to OVA_CytoT_ Cat-PAMs & CpG exhibited increased MHC II expression over all other vaccine formulations assessed. Overall, these data suggest unmodified CpG generally enhanced BMDC CD86 and MHC II cell surface expression relative to lipidated CpG when supplementing both peptide and PAM formulations. Specifically, statistical significance over 5’-lipidation was observed for all but OVA­_CytoT_ Z-PAMs whereas a significant increase over 3’-lipidation was only seen for MHC II expression in cells treated with PAMs. In light of these properties, unmodified CpG was motivated forward for further experimentation.

### OVA_CytoT_ PAM\CpG Association

In order to determine whether electrostatic complexation between CpG and OVA_CytoT_ PAMs had been achieved, further physical characterization was necessary. Förster Resonance Energy Transfer (FRET) was employed to provide evidence of micelle|CpG affiliation (21, 66). FRET studies were conducted with TAMRA entrapped OVA_CytoT_ PAMs and FAM-CpG, with single fluorophore formulations used as baseline spectral behavior with the donor peak found at ∼ 525 nm (FAM emission wavelength) and the acceptor peak observed at ∼ 580 nm (TAMRA emission wavelength) (**Figure 3a**). When fluorophore associated OVA_CytoT_ PAMs and CpG were incubated together, a large decrease in donor fluorescence was found with a slight increase in acceptor fluorescence consistent with close physical proximity of the two species. Interestingly, this was observed for both OVA_CytoT_ Cat-PAMs and OVA_CytoT_ Z-PAMs.

**Figure 3.**
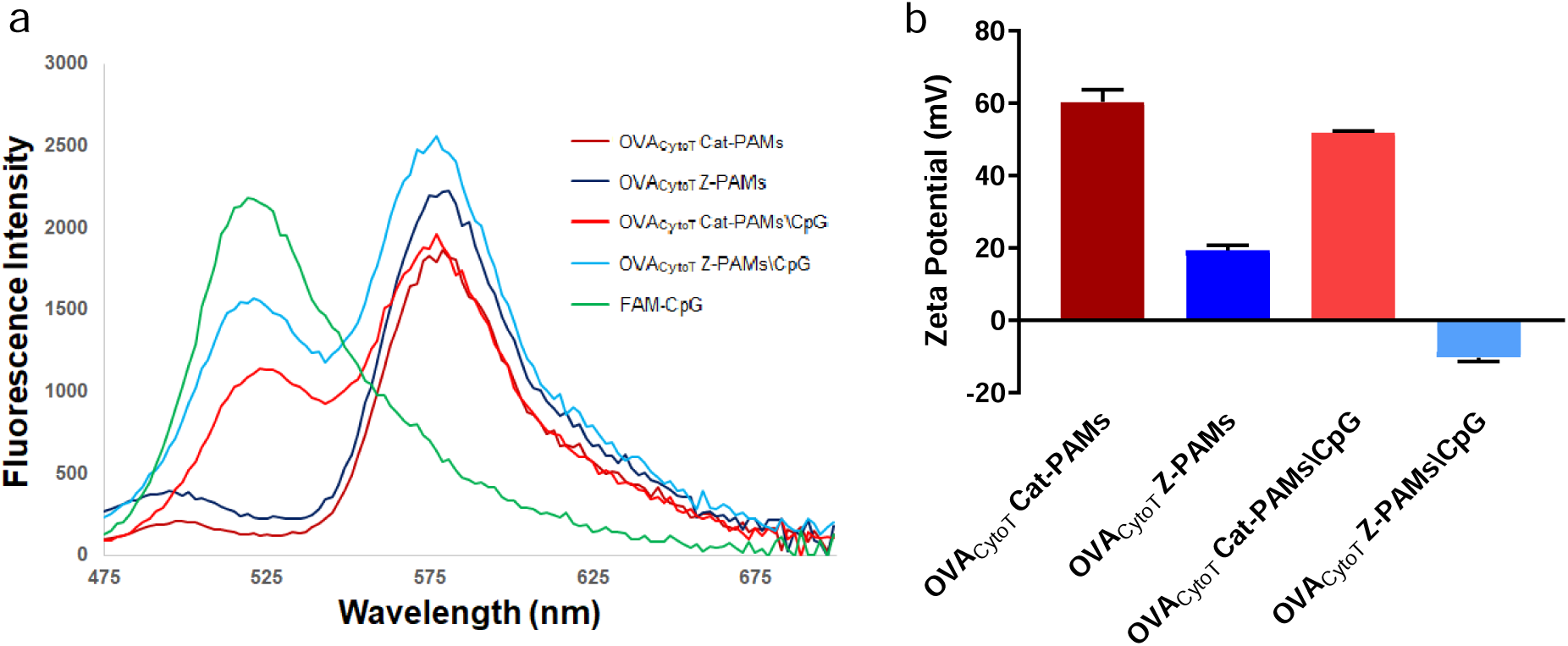
CpG readily associates with OVA_CytoT_ PAMs. **a** Fluorescence spectra data from a Förster Resonance Energy Transfer (FRET) study show the presence of only the donor peak (∼ 525 nm) for FAM-CpG and the acceptor peak (∼ 580 nm) for TAMRA entrapped in either of the OVA_CytoT_ PAM formulations. When OVA_CytoT_ PAMs and CpG were mixed during fabrication, the resulting formulations (*i.e.,* OVA_CytoT_ Cat-PAMs\CpG and OVA_CytoT_ Z-PAMs\CpG) underwent electrostatic complexation as demonstrated by a considerable reduction and small enhancement in donor and acceptor fluorescence, respectively. **b** Zeta potential of OVA_CytoT_ PAMs unsurprisingly showed OVA_CytoT_ Cat-PAMs were more positively charged than OVA_CytoT_ Z-PAMs. When CpG was present, the observed zeta potential decreased for both formulations though no evaluation for statistical significance was performed.

To complement the FRET studies, zeta potential measurements were collected to probe the surface charge of these OVA_CytoT_ PAM\CpG complexes in PBS (**Figure 3b**). OVA_CytoT_ Cat-PAMs were found to have a relatively large, positive zeta potential (+60.3 mV) indicative of a highly cationic particle surface. Likewise, OVA_CytoT_ Z-PAMs were observed to have a modestly, cationic surface (*i.e.*, zeta potential of +19.4 mV). When OVA_CytoT_ Cat-PAMs were fabricated with negatively charged CpG, both resulting formulations exhibited a lower surface potential (*i.e.*, +51.8 mV and −10.2 mV, respectively) implying a reduction in their surface charges. It is worth noting OVA_CytoT_ Cat-PAMs with and without CpG were near the detection limit of the instrument (± 60 mV) and accordingly, no statistical analyses were conducted for these data.

### *In vivo* assessment of OVA_CytoT_ PAMs\CpG

Having fabricated adjuvant complexed micelles with various electrostatic properties, *in vivo* studies assessing their ability to induce strong cell-mediated immune responses against their incorporated cytotoxic T cell antigen were performed. Following two rounds of immunization at days 0 and 10 with micelle formulations and peptide controls according to **Table III**, sex-matched C57BL/6J mice were euthanized at day 20 and their lymph nodes and spleens harvested for *ex vivo* and *in vitro* analysis. Flow cytometry was used to quantify the Ag-specific cytotoxic T cell population present in the collected secondary lymphoid tissues using a FITC CD8 antibody and an MHC I haplotype matched APC SIINFEKL tetramer. After gating for live cells on forward scatter versus side scatter plots, the CD8^+^ lymphocytes (*i.e.,* FITC fluorescent intensity > 25) were selected. These cells were then further gated in the APC channel utilizing appropriate stain controls to decipher signal versus background allowing for the identification of SIINFEKL-specific cytotoxic T cells in the draining lymph nodes (**Figure S2**). These data indicated exciting differences among the vaccine groups (**Figure 4a**). Animals vaccinated with OVA_CytoT_ peptide with or without CpG supplementation showed limited, statistically insignificant, increases in antigen-specific CD8^+^ lymphocytes (0.059% and 0.049%) as compared to the PBS vaccinated control group (0.035%). Similar results were observed with mice vaccinated with OVA_CytoT_ Cat-PAMs (0.060%) or OVA_CytoT_ Z-PAMs (0.061%).

**Figure 4.**
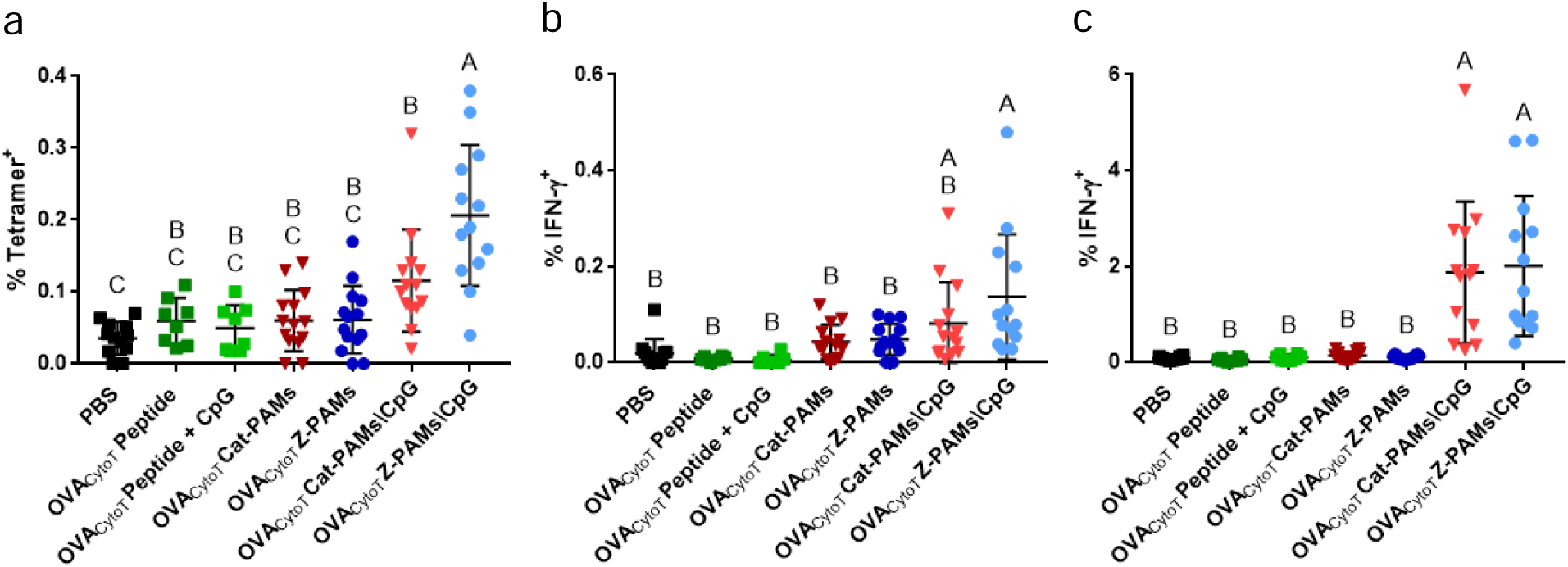
CpG complexed micelles induce enhanced cytotoxic T cell immune responses. Vaccine formulation affects the generation of **an** antigen-specific draining lymph node CD8+ T cells and antigen responsive CD8+ T cells in the **b** draining lymph nodes and **c** spleen. Specifically, SIINFEKL Tetramer^+^ and IFN-γ^+^ CD8^+^ T cell induction was dependent on co-delivery of peptide antigen and adjuvant confined to micelles. Within each graph, groups that possessed different letters have statistically significant differences in mean (p ≤ 0.05) whereas those that have the same letter do not.

In contrast, mice that received CpG-complexed OVA_CytoT_ PAMs showed considerable improvement in the presence of SIINFEKL-specific cytotoxic T cells with highly positively charged micelles (*i.e.,* OVA_CytoT_ Cat-PAMs\CpG) inducing a weaker response (0.115% versus 0.206%) than slightly negatively charged micelles (*i.e.,* OVA_CytoT_ Z-PAMs\CpG). Spleen data for SIINFEKL tetramer were inconclusive due to significant cell autofluorescence observed in the APC and FITC channels (*data not shown*).(69–71)

The magnitude of the vaccine-induced immune response was also tested by exploring lymphocyte antigen restimulatory capacity *in vitro*. When cytotoxic T cells recognize their cognate antigen in the context of MHC I and a costimulatory signal, they activate and produce the cytokine IFN-γ. Exposing cells to a protein transport inhibitor (*i.e.*, Brefeldin A) together with stimulation prevents them from secreting cytokines allowing for their subsequent detection with antibody-based staining and flow cytometry analyses. Single cell suspensions isolated from draining lymph nodes were pulsed with OVA_CytoT_ peptide for 6 hours, after which cells were stained with FITC CD8 antibody and APC IFN-γ antibody. After gating for live cells on forward scatter versus side scatter plots, CD8^+^ lymphocytes (*i.e.,* FITC fluorescent intensity > 25) were selected. These cells were then further gated for APC using appropriate stain controls to determine signal versus background and quantification of the fraction of IFN-γ^+^ cytotoxic T cells in the draining lymph nodes (**Figure S3**). Flow cytometry data compiled from all analyzed samples showed exciting differences between select vaccine groups (**Figure 4b**). Animals vaccinated with OVA_CytoT_ peptide both with and without CpG supplementation were statistically indistinguishable for their proportion of IFN-γ^+^ CD8^+^ lymphocytes (0.006% and 0.008%) from the PBS vaccinated control (*i.e.,* 0.019%). Mice that received the micelle formulations alone (*i.e.,* OVA_CytoT_ Cat-PAMs and OVA_CytoT_ Z-PAMs) were found to have lymph node and spleen Ag specific CD8^+^ cell populations statistically indistinguishable from those of the PBS treated control (p ≤ 0.05). CpG-complexed OVA_CytoT_ PAMs (*i.e.,* OVA_CytoT_ Cat-PAMs\CpG and OVA_CytoT_ Z-PAMs\CpG) induced more activated on-target cells (*i.e.*, 0.081% and 0.137%) with statistical significance observed for the zwitterionic micelle formulation.

While autofluorescence was found to be a complication with analyzing SIINFEKL specific T cell receptor (TCR) expression on splenocytes, no such issue was observed with the APC IFN-γ antibody. Similar to lymph node assessment, single cell suspensions isolated from spleens were pulsed with OVA_CytoT_ peptide and brefeldin for 6 hours, after which they were stained with FITC CD8 antibody and APC IFN-γ antibody. After gating for live cells on forward scatter versus side scatter plots, the CD8^+^ T cell sub-population (*i.e.,* FITC fluorescent intensity > 25) was selected. These cells were then further gated for APC utilizing appropriate stain controls to decipher signal versus background allowing for the identification of IFN-γ producing lymphocytes in the spleen (**Figure S4**). Flow cytometry data compiled from all analyzed these samples also showed exciting differences between the vaccines groups (**Figure 4c**). Again, animals vaccinated with OVA_CytoT_ peptide with or without CpG supplementation or OVA_CytoT_ PAMs exhibited antigen-specific CD8^+^ lymphocyte proportions (0.05% and 0.14%) statistically indistinguishable from PBS vaccinated controls (0.09%). Alternatively, mice that received CpG-complexed OVA_CytoT_ PAMs showed considerable improvement in IFN-γ producing cytotoxic T cells with both highly positively charged micelles (*i.e.*, OVA_CytoT_ Cat-PAMs\CpG) and slightly negatively charged micelles (*i.e.*, OVA_CytoT_ Z-PAMs\CpG) inducing strong CD8^+^ T cell responses of 1.88% and 2.01%, respectively. Gating for CD8^-^ lymphocytes and splenocytes showed no significant increase of IFN-γ^+^ cells when PAMs and CpG were co-delivered (**Figure S5**, **Figure S6**, and **Figure S7**) indicating CpG complexation does not contribute to additional CD8^-^ lymphocyte IFN-γ production.

### Evaluation of OVA_BT_ PAMs with CpG

Having observed that OVA_CytoT_\CpG micellar co-delivery enhanced cell-mediated immune responses, the interchangeability of this observation was assessed by employing a different bioactive peptide, a linked recognition B cell and helper T cell antigen (OVA_BT_). Due to the role surface charge played in OVA_CytoT_ PAM vaccination (**Figure 4**) and this phenomenon having been previously established with OVA_BT_ (20), a new experimental group was created to charge match micelles and CpG, for which calculations determined a 16:1 Z-PA:Cat-PA ratio was needed. Indeed, mixed micelle formulations (OVA_CytoT_ Cat/Z-PAMs (+26.6 mV) and OVA_CytoT_ Cat/Z-PAMs\CpG (−9.0 mV)) manifested comparable charge behavior to their zwitterion-like counterparts (OVA_CytoT_ Z-PAMs (+19.4 mV) and OVA_CytoT_ Z-PAMs\CpG (−10.2 mV)) (**Figure 5a**). Similar data were collected for OVA_BT_ PAM formulations (**Figure 5b**), though CpG complexed micelles remained modestly positively charged in contrast to their OVA_CytoT_ PAM analogues (+ 40.7 mV versus +38.0 mV and +14.7 mV versus +11.8 mV, respectively). FRET results indicated cationic (*i.e.,* OVA_BT_ Cat-PAMs) and charge matched (*i.e.,* OVA_BT_ Cat\Z-PAMs) products achieved CpG association as seen by significant donor fluorescence decrease and slight acceptor fluorescence increase (**Figure S8**). In sharp contrast to OVA_CytoT_ Z-PAMs, OVA_BT_ Z-PAMs did not appear to readily associate with CpG as evidenced by a limited change observed between their fluorescence spectra (**Figure S8**). This uniquely provides an opportunity to assess the importance of CpG complexation in micellar immunogenicity through the comparison of OVA_BT_ Z-PAMs + CpG and OVA_BT_ Cat/Z-PAMs\CpG.

**Figure 5.**
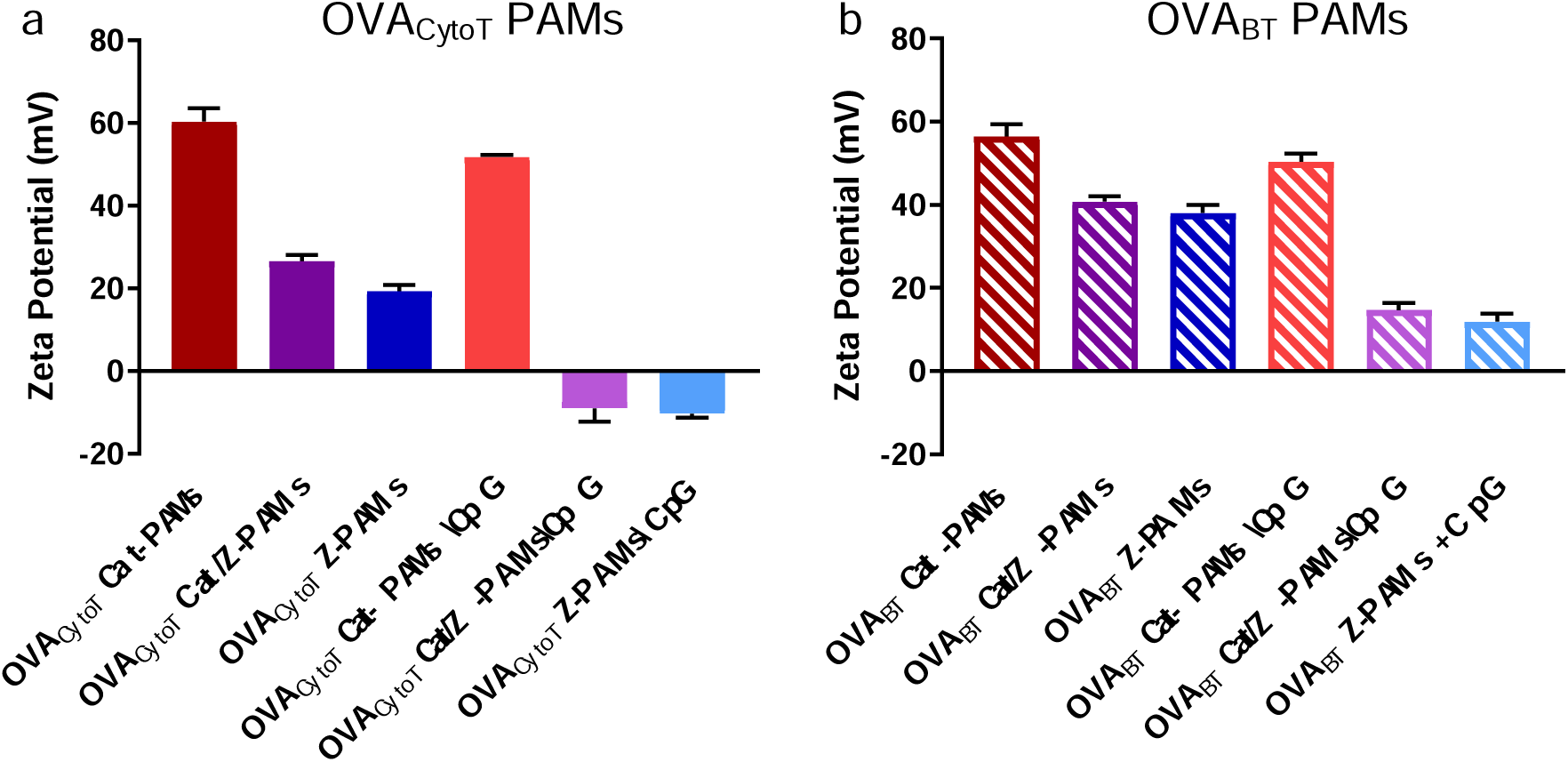
Peptide and CpG presence alter OVA PAM charge. Zeta potential was measured for different micelle formulations incorporating **a** OVA_CytoT_ and **b** OVA_BT_. As predicted based on chemical structure, Cat-PAMs were more positively charged than Z-PAMs. Cat/Z groups indicate a mixed PA population was used to form the PAMs (specifically a 16:1, Z to Cat ratio). Whereas the inclusion of CpG caused a modest reduction of charge in Cat-PAMs, Z-PAMs saw a more drastic change.

### *In vitro* assessment of OVA_BT_ PAMs with CpG

In order to assess the influence CpG complexation has on the biological interactions of OVA_BT_ PAMs, *in vitro* cellular studies were conducted. Different formulations were incubated with either phagocytic cells (*i.e.,* MØs) or non-phagocytic cells (*i.e.,* T cells) for 1 hour, after which fluorophore-labelled CpG association with cells was evaluated by flow cytometry (**Figures 6a - 6b**) to explore uptake specificity. Results indicated that OVA_BT_ Cat-PAMs\CpG have the strongest cell association (MFI - 375) of the tested cell types likely due to their considerable positive charge helping to facilitate non-specific cell association in agreement with our previously published results for OVA_BT_ Cat-PAMs alone (20). OVA_BT_ Cat/Z-PAMs facilitated a modest increase in CpG internalization (MFI - 182) by MØs over CpG alone (MFI - 120) without a significant increase in T cell internalization (MFI - 3.90 versus MFI - 2.56). These results were similar to OVA_BT_ Z-PAMs with non-complexed CpG indicating the presence of micelles alone may facilitate CpG internalization enhancement for modestly charged formulations. Next, the capacity for this enhanced internalization behavior to influence CpG bioactivity was also explored. BMDCs exposed to different stimuli were evaluated for their activation state by flow cytometric analysis of three different cell surface receptors (*i.e.,* CD40, CD86, and MHC II) (**Figures 6c - 6e**). Results indicate that the presence of CpG (alone, complexed, or non-complexed) is the primary facilitator of BMDC activation with every CpG containing group enhancing expression over the PBS control. A modest additional enhancement in the expression of costimulatory markers (CD40 at MFI - 66.8 and CD86 at MFI - 2220) over CpG control (CD40 at MFI - 53.9 and CD86 at MFI - 1710) was seen with OVA_BT_ Cat-PAMs likely due to their increased cell association capacity.

**Figure 6.**
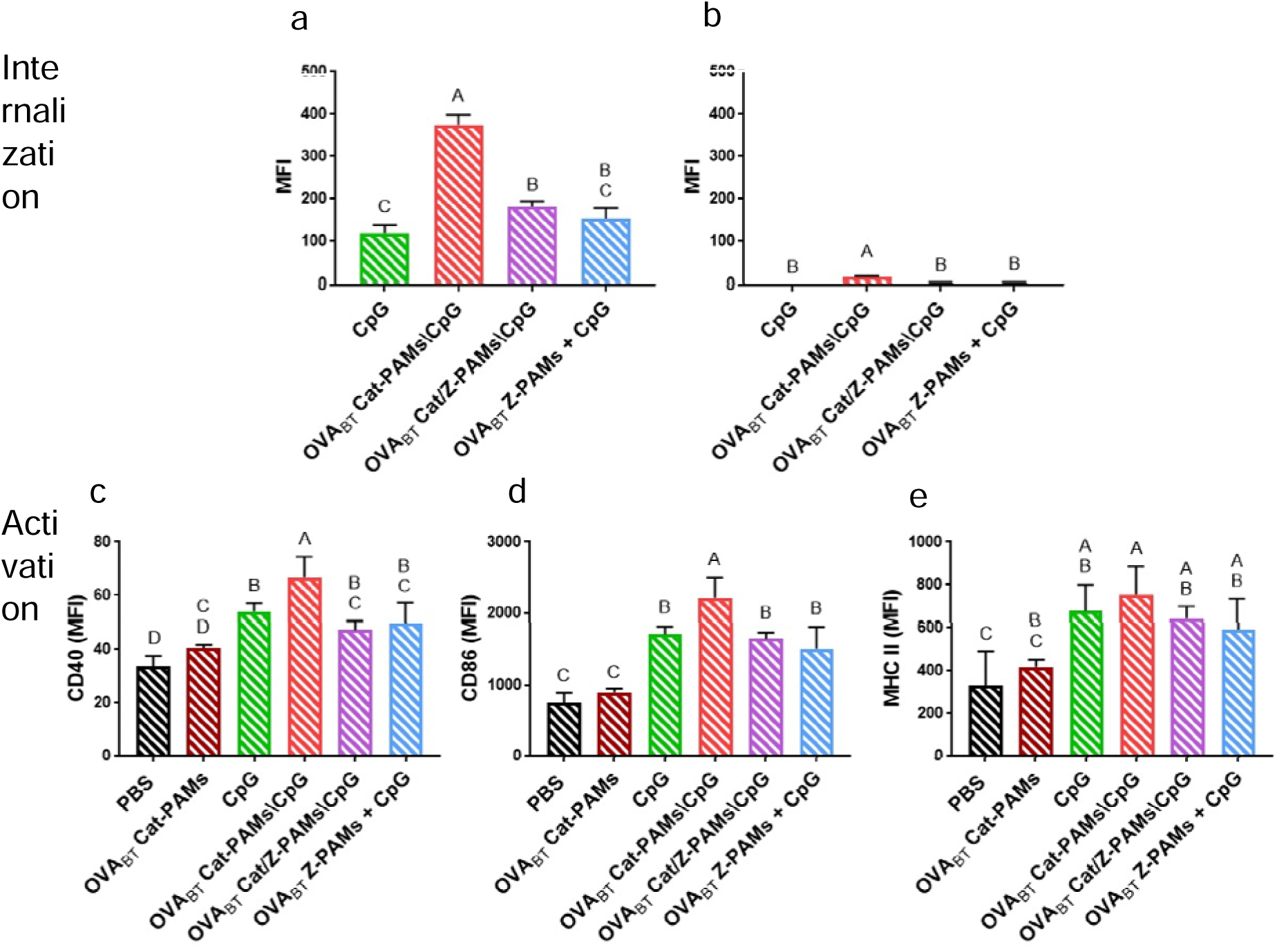
PAM charge influences CpG-cell interactions. **a** Macrophages or **b** T cells were incubated with indicated formulations for 1 hour, before being fixed and evaluated by flow cytometry. OVA_BT_ Cat-PAMs increased CpG cell association for phagocytic cells (*i.e.*, MØs) as well as non-phagocytic cells (*i.e.,* T cells) though not as significantly for the latter population. BMDCs were stimulated with various vaccine formulations for 24 hours after which activation was assessed by the presence of surface cell markers **c** CD40, **d** CD86 and **e** MHC II. BMDC surface markers for OVA_BT_ Cat-PAMs were statistically indistinguishable from PBS control. In contrast, exposure to CpG significantly increased BMDC activation marker expression regardless of which formulation it was included. The complexation of CpG with OVA_BT_ Cat-PAMs did not alter MHC II presentation but did further enhance CD40 and CD86 expression. Within a graph, groups that possess different letters have statistically significant differences in mean (*p* ≤ 0.05) whereas those that possess the same letter do not.

### *In vivo* assessment of OVA_BT_ PAMs with CpG

Having assessed the role of CpG complexation on *in vitro* BMDC activation, *in vivo* studies were conducted to determine its impact on antigen-specific immune responses. Mice were immunized subcutaneously in the nape of the neck each with one of the formulations outlined in **Table IV** at weeks 0 and 4. The presence of OVA_BT_-specific IgG antibody was evaluated by ELISA using serum collected 2 weeks post-boost vaccination (*i.e.*, week 6) (**Figure 7**). The OVA_BT_ Cat/Z-PAMs\CpG and OVA_BT_ Z-PAMs + CpG formulations (mean IgG titers of 158,000 and 172,000, respectively) induced total antigen specific IgG production (**Figure 7a**) an order of magnitude greater than the two other vaccine formulations (*i.e.*, peptide + CpG and OVA_BT_ Cat-PAMs/CpG which had mean IgG titers of 5,380 and 16,600, respectively). IgG subtypes (*i.e.*, IgG1, IgG2a, and IgG3) evaluated at the same time point (**Figure 7b - 7d**) all showed similar trends as total IgG indicating micelle charge (*i.e.,* mildly cationic versus highly cationic) influenced B cell associated immune responses more than CpG complexation.

**Figure 7.**
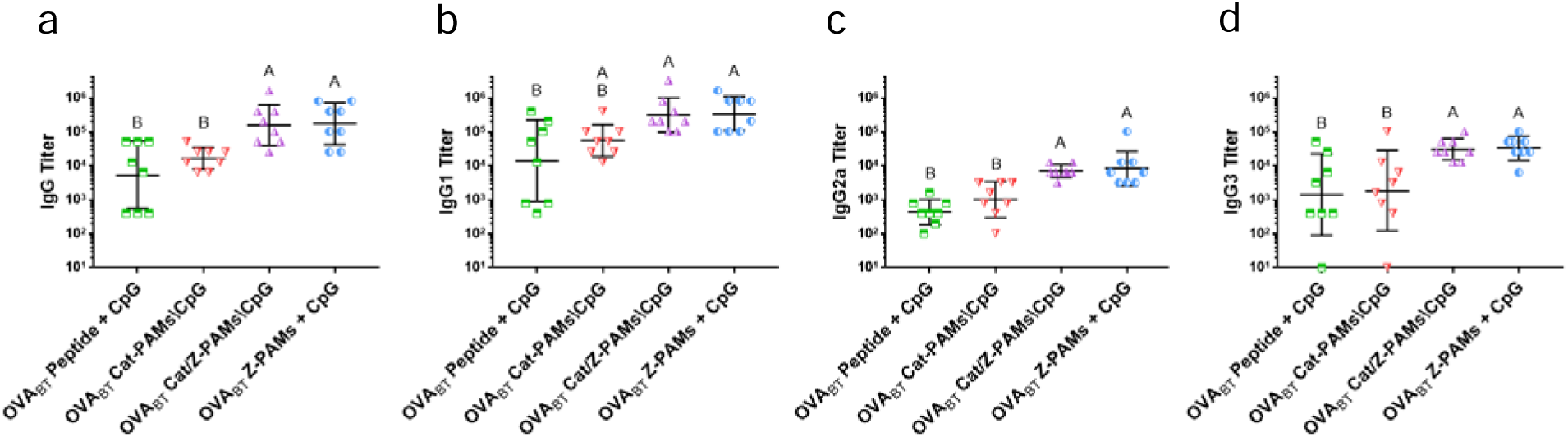
Micelle charge, not electrostatic antigen\adjuvant co-localization, correlates with antibody production. Serum was collected from mice injected with the indicated vaccine formulations two weeks post-boost immunization and analyzed for OVA_BT_-specific antibody content by ELISA. **a** Total IgG antibody production results revealed moderately positively charged PAMs (*i.e.*, Cat/Z-PAMs\CpG and Z-PAMs + CpG) induced higher antibody production than adjuvant supplemented peptide (*i.e.*, peptide + CpG) and highly positively charged PAMs (*i.e.*, Cat-PAMs\CpG). Three major IgG subtypes - **b** IgG1, **c** IgG2a, and **d** IgG3 - showed similar trends as IgG with IgG1 as the dominant subtype followed by more modest production of IgG2a and IgG3. Within a graph, groups that possess different letters have statistically significant differences in mean (*p* ≤ 0.05) whereas those that possess the same letter do not.

In addition to antibody production, cell-mediated responses are a vital facet of vaccination. One aspect of the adjuvanticity of CpG is its ability to facilitate the development of antigen specific lymphocytes that respond rapidly to antigen re-stimulation. To assess this behavior, single cell suspensions of lymphocytes (**Figures 8a - 8c**) and splenocytes (**Figures 8d - 8f**) harvested from vaccinated animals were stimulated with OVA_BT_ antigen *in vitro*. After 24 hours, the concentration of supernatant cytokines (*i.e.,* IL-2, IFN-γ, and TNF-α) were measured by ELISA. The results indicated that both cell populations responded quite similarly with the marked increases in cytokine production expected from memory cells upon restimulation only observed for formulations where the antigen and adjuvant were complexed together (*i.e.,* OVA_BT_ Cat-PAMs/CpG and OVA_BT_ Cat/Z-PAMs/CpG).

**Figure 8.**
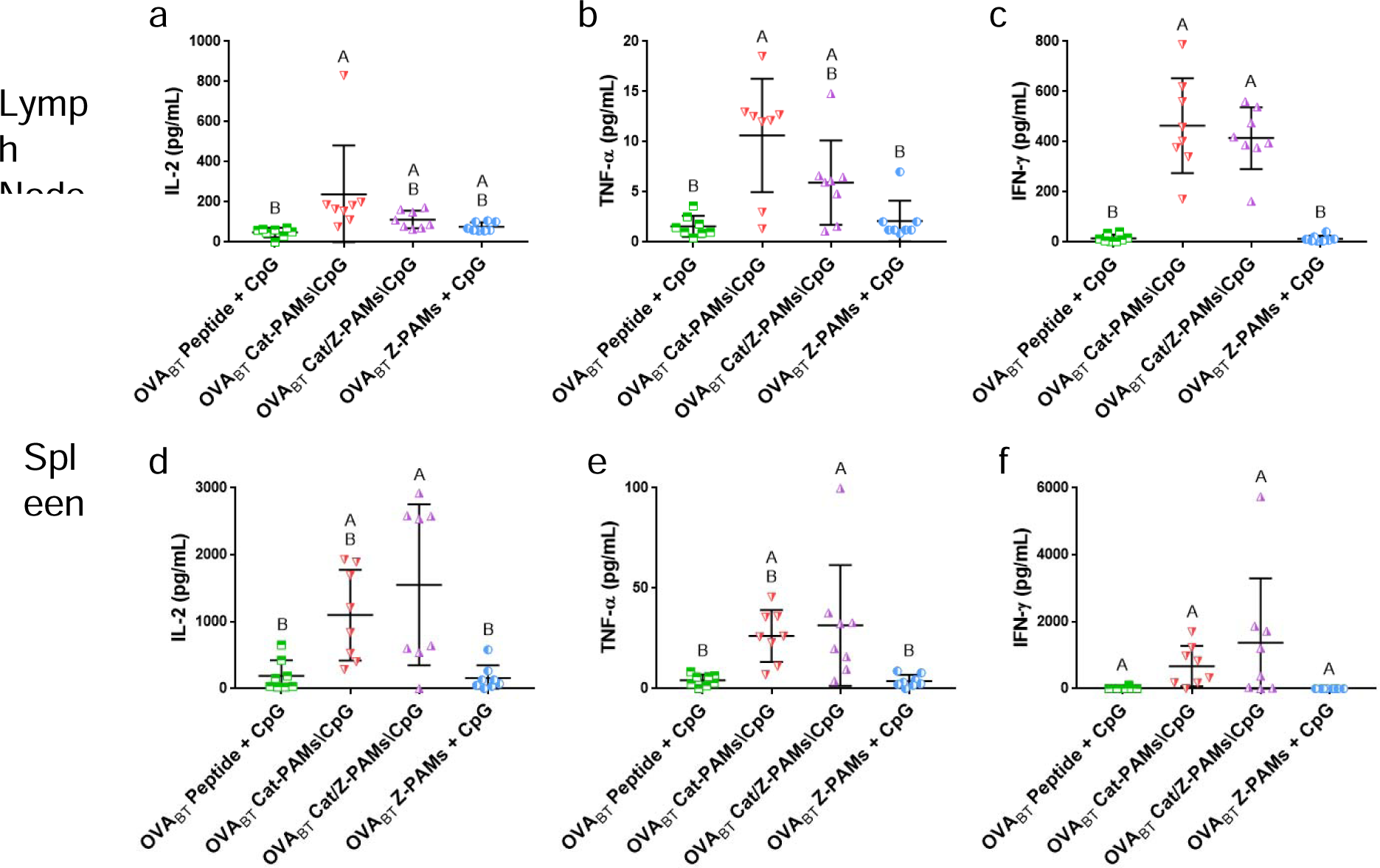
Electrostatic antigen\adjuvant co-localization affects lymphocyte antigen responsiveness. Cell culture supernatants collected from OVA_BT_ peptide stimulated **a - c** lymphocytes and **d - f** splenocytes 72 hours after re-stimulation with OVA_BT_ peptide were analyzed for their cytokine content (*i.e.*, IL-2, IFN-γ, and TNF-α) by ELISA. Both lymph node cells and spleen cells possessed a similar trend where antigen\adjuvant complexed formulations (*i.e.*, Cat-PAMs\CpG and Cat/Z-PAMs\CpG) induced appreciable cytokine production over vaccines where the antigen and adjuvant were not closely associated (*i.e.*, peptide + CpG and Z-PAMs + CpG). Within a graph, groups that possess different letters have statistically significant differences in mean (*p* ≤ 0.05) whereas those that possess the same letter do not.

## Discussion

While PAMs have shown significant potential as vaccine delivery systems (21,62), it is still not well understood how their design parameters (*e.g.*, peptide charge block / surface charge, adjuvant association, degree of lipidation, *etc.*) should be tuned to induce the most optimal host immune responses. Our recent efforts have indicated that PAM size and surface charge can be readily altered by manipulating their charged block region which directly enhances or suppresses host immune responses to incorporated peptide antigens (22,23). Considering this observation, TEM and zeta potential studies were performed for each vaccine formulation to assess their shape, size, and surface charge. No perceivable size or shape differences were observed upon CpG complexation for any of the formulations studied (**Figure 1** and **Figure S1**) and each OVA_CytoT­_ containing formulation was in the size range (10 - 100 nm) which has been found to correlate with improved immunogenicity of the incorporated peptide antigen (20,72–74).

As adjuvant incorporation with PAM vaccines is non-trivial, both electrostatic complexation and hydrophobic association were considered as possible means for colocalizing PAs and adjuvants. To evaluate the impact incorporation method has on adjuvant bioactivity, electrostatically complexed and hydrophobically associated vaccine formulations containing a cytotoxic T cell epitope were assessed for their capacity to activate BMDCs *in vitro* (**Figure 2**). Interestingly, electrostatic complexation was found to better induce the upregulation of BMDC surface markers (*i.e.*, CD86 and MHC II) over both CpG along and PAMs with hydrophobically associated CpG making it the more bioactive method for adjuvant|PAM colocalization. While 5’- and 3’-lipidated CpG variants both resulted in less optimal cell surface marker enhancement, subtle differences were seen between the two lipidation schemes suggesting caution should be taken when decisions are made regarding covalent conjugation of a bioactive molecule like CpG (67). To verify adjuvant|PAM complexation indeed occurred as hypothesized, *in vitro* FRET analysis was performed (**Figure 3**). Specifically, this study (**Figure 3a**) showed a larger reduction in donor signal for OVA_CytoT­_ Cat-PAMs\CpG PAMs than OVA­_CytoT_ Z-PAMs\CpG PAMs suggesting that there may be more FAM-CpG in proximity to the TAMRA trapped within the former over the latter. Furthermore, when zeta potential analysis was performed (**Figure 3b**), a distinct reduction in charge upon addition of CpG was observed indicating association of the negatively charged CpG to the micelle surface. In concert, these experiments provided evidence that both OVA_CytoT_ Cat-PAMs and OVA_CytoT_ Z-PAMs are capable of CpG electrostatic complexation.

Intriguingly, when verifying whether electrostatic complexation was achieved between the OVA_BT_ PAMs formulations and CpG, FRET data suggested OVA­_BT­_ Z-PAMs did not successfully complex CpG whereas the OVA­_BT­_ Cat-PAMs and OVA_BT_ Cat/Z-PAMs formulations did (**Figure S8**). This outlier highlights the notion that zeta potential alone is not the only factor that governs electrostatic complexation (**Figure 5**) while also providing the opportunity to evaluate whether CpG co-delivery with the antigen was necessary to enhance immunogenicity or if co-treatment by itself is sufficient. Whereas the OVA_CytoT_ studies suggested the CpG incorporation method mattered (**Figure 2**), the OVA­_BT­_ PAM *in vitro* stimulation studies (**Figure 6c-e**) revealed no significant differences in cell surface marker expression for BMDCs incubated with OVA­_BT_­ PAMs bearing electrostatically complexed CpG (*i.e.*, OVA_BT_ Cat/Z-PAMs\CpG) or OVA­_BT­_ PAMs possessing similar surface charge (**Figure 5b**) co-treated with non-complexed CpG (*i.e.*, OVA­_BT­_ Z-PAMs + CpG). It appeared that the presence of unmodified CpG alone was sufficient to drive the upregulation of co-stimulatory marker expression as both of these formulations were not only similar to each other but also the positive control of CpG alone. In contrast, the more cationic OVA_BT_ PAM formulation (*i.e.*, OVA_BT_ Cat-PAMs\CpG - **Figure 5b**) was found to be more readily internalized by APCs (**Figure 6a**) which likely explains the enhancement in CD40 and CD86 expression observed in BMDCs (**Figure 6c-d**) though no subsequent improvement in MHC II expression was observed (**Figure 6e**). As these molecules play an important role in the activation of adaptive immune responses, adjuvant incorporation method likely impacts vaccine formulation incorporating B and helper T cell epitopes as well as those containing a cytotoxic T cell epitope.

Flow cytometry and ELISA analysis generally corroborated the importance of CpG but also suggested surface charge (as measured by zeta potential) plays an important role in inducing productive immune responses. While electrostatic complexation of CpG on OVA_CytoT_ PAMs generally enhanced the relative proportion of lymph node-derived, antigen specific or activated CD8+ T cells (**Figure 4a-b**), statistical significance over their respective, bare PAMs controls was only achieved in mice vaccinated with the moderately charged, zwitterion-like formulation (*i.e.*, OVA_CytoT_ Z-PAMs\CpG). Interestingly, the more cationic formulation (*i.e.*, OVA_CytoT_ Cat-PAMs\CpG) was found to induce stronger BMDC activation (based on MFI of CD86 and MHC II - **Figure 2**) *in vitro* than its moderately charged, zwitterion-like counterparts. The difference between these results is likely due to the highly complex environment found *in vivo* where cationic micelles will more indiscriminately associate with any cells and limit trafficking to the draining lymph node as we have previously observed with OVA_BT_ Cat-PAMs (20). Consistent with previous studies wherein CpG conjugation to both uncharged polymeric nanoparticle (75,76) and cationic gelatin-based nanoparticle vaccines (77) have been shown to facilitate antigen specific activation of T cells, we observed CpG complexation to PAM formulations induced a significant increase in IFN-γ+ CD8+ T cell production in the spleen (relative to the formulations without a delivery vehicle), which was found to be insensitive to the charge of the particle (**Figure 4c**).

Similarly, more moderately charged, CpG-complexed OVA_BT_ micelles (*i.e.*, OVA_BT_ Cat/Z-PAMs\CpG) induced higher antibody titers (**Figure 7**) than the analogous highly cationic formulation (*i.e.*, OVA_BT_ Cat-PAMs\CpG). Interestingly, it was also determined that CpG complexation was not required to include a robust immune response as similar high antibody titers were observed with soluble CpG-supplemented OVA_BT_ micelles (*i.e.*, OVA_BT_ Z-PAMs + CpG) as were seen with OVA_BT_ Cat/Z-PAMs\CpG (**Figure 7**). In contrast, CpG complexation to micelles did appear to be necessary to produce lymphocytes and splenocytes capable of cytokine production upon antigen restimulation *in vitro* (**Figure 8**). When combined with the companion APC stimulation *in vitro* data (**Figure 6**) and previous lymph node trafficking data (20), it appears likely that cationic micelles limit the capacity for the full antigen load to be appropriately delivered to secondary lymphoid tissues. Moreover, despite PAM\CpG complexation appearing to enhance cytokine expression, PAM + CpG co-delivery appeared sufficient for producing high antibody titers.

## Conclusion

Taken together, this research demonstrated that PAMs are a multifunctional immunostimulatory delivery platform capable of inducing immune responses against micelle-incorporated cytotoxic T cell peptide antigen or linked recognition B-T cell peptide antigen complementing previous work that showed their capacity to enhance cell-mediated and antibody-mediated immune responses (17–21,62,63). After providing evidence of successful micellization of triblock OVA_CytoT_ PAs similarly to our previous OVA_BT_ PA work as well as facilitating association of both of these systems with CpG, *ex vivo* and *in vitro* studies showed that PAM\CpG vaccines could successfully induce desirable immune responses *in vivo*. The CpG adjuvant incorporation method was found to be extremely valuable to the immunogenicity of the PAMs for either arm of the immune response induced. Micelle charge was found to almost universally play a direct role in the strength of the corresponding downstream immune response whereas complexation was necessary to maximize cell-based stimulatory and restimulatory effects. Overall, this research provides a strong foundation that PAM charge and adjuvant complexation can be tuned to achieve complex, desirable immune response induction. While promising, future work is needed to further optimize PAM immunogenicity for this application specifically focusing on the finer tuning of PAM charge as well as the incorporation of various additional factors such as other adjuvants, DNA, and/or siRNA to leverage these materials for disease-specific vaccine applications.

## Acknowledgements

We thank the NIH Tetramer Core Facility at Emory University for Providing Tetramers. The SIINFEKL Tetramer was funded by the NIH Grant R01GM103841 (PI - Adam G. Schrum). This research was supported by University of Missouri Start-up Funds, the University of Missouri Research Council, and the PhRMA Foundation.

