## Supplementary Information for "Adjuvant Delivery Method and Nanoparticle Charge Influence Peptide Amphiphile Micelle Vaccine Bioactivity"

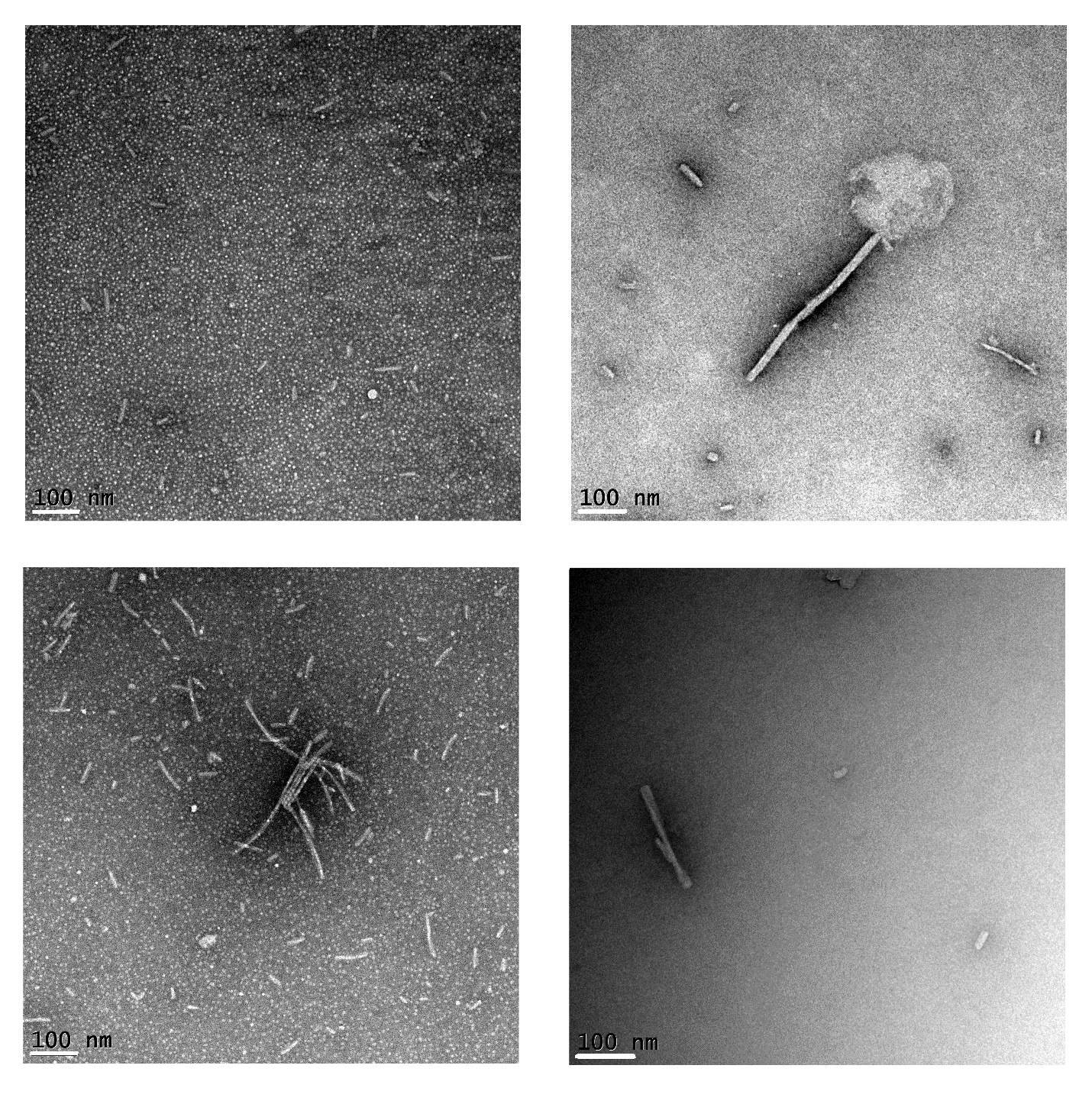

d

c

b

a

**Figure S1.** Transmission electron microscopy (TEM) analysis shows the inclusion of Lipid-CpG does not significantly alter nanoparticle formation. Specifically, the presence of **a** and **b** Lipid-CpG 5’ or **c** and **d** Lipid-CpG 3’ yields no major changes in micellar nanoarchitecture with **a** and **c** OVA_CytoT_ Cat-PAs forming spherical and short cylindrical micelles and **b** and **d** OVA_CytoT_ Z-PAs self-assembling into long cylindrical micelles. A 10 μM PA concentration and a scale bar of 100 nm were used for all micrographs.

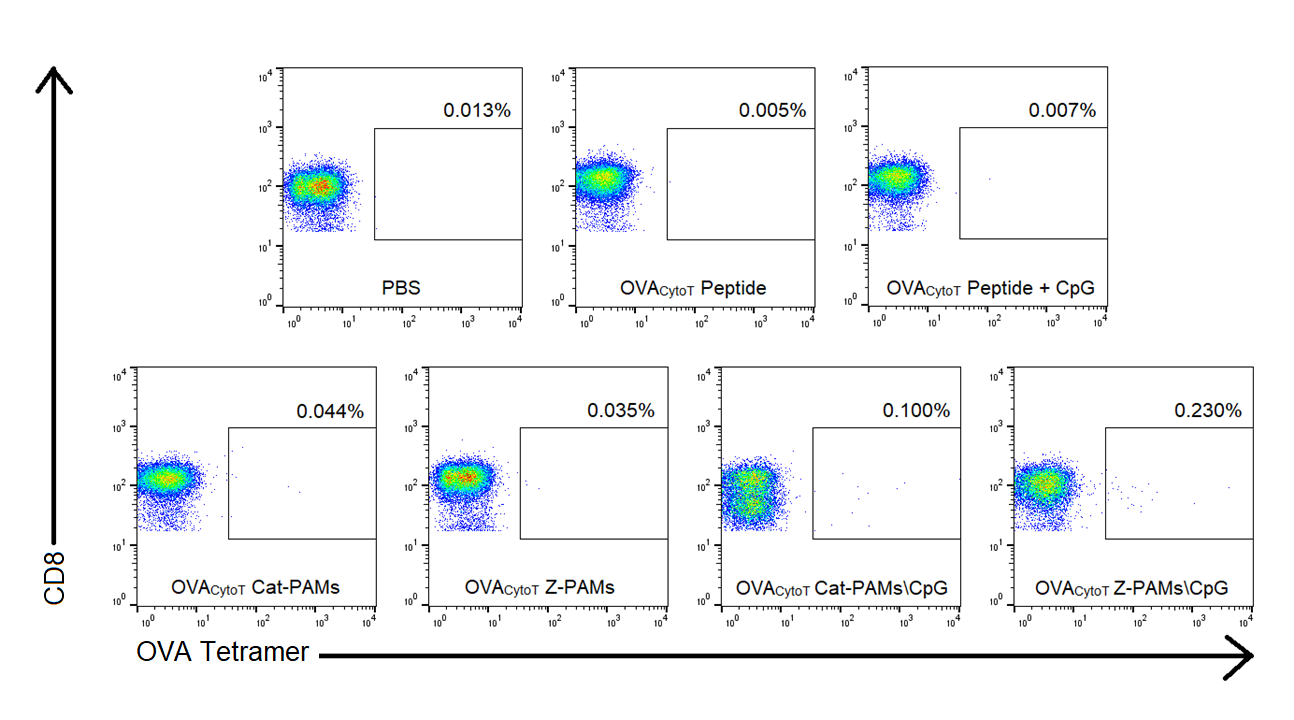

**Figure S2.** Representative lymph node CD8 versus SIINFEKL Tetramer flow cytometry dot plots. Population gating was compensated by utilizing no stain and single stain controls after which the data was analyzed by Flo Jo software. The presence of cell surface CD8 was first gated to separate out the cytotoxic T cell population from other living lymphocytes. This population was then investigated for the presence of cell surface associated OVA peptide specific T cell receptor (TCR) to determine the percentage of SIINFEKL Tetramer^+^ CD8^+^ T cells.

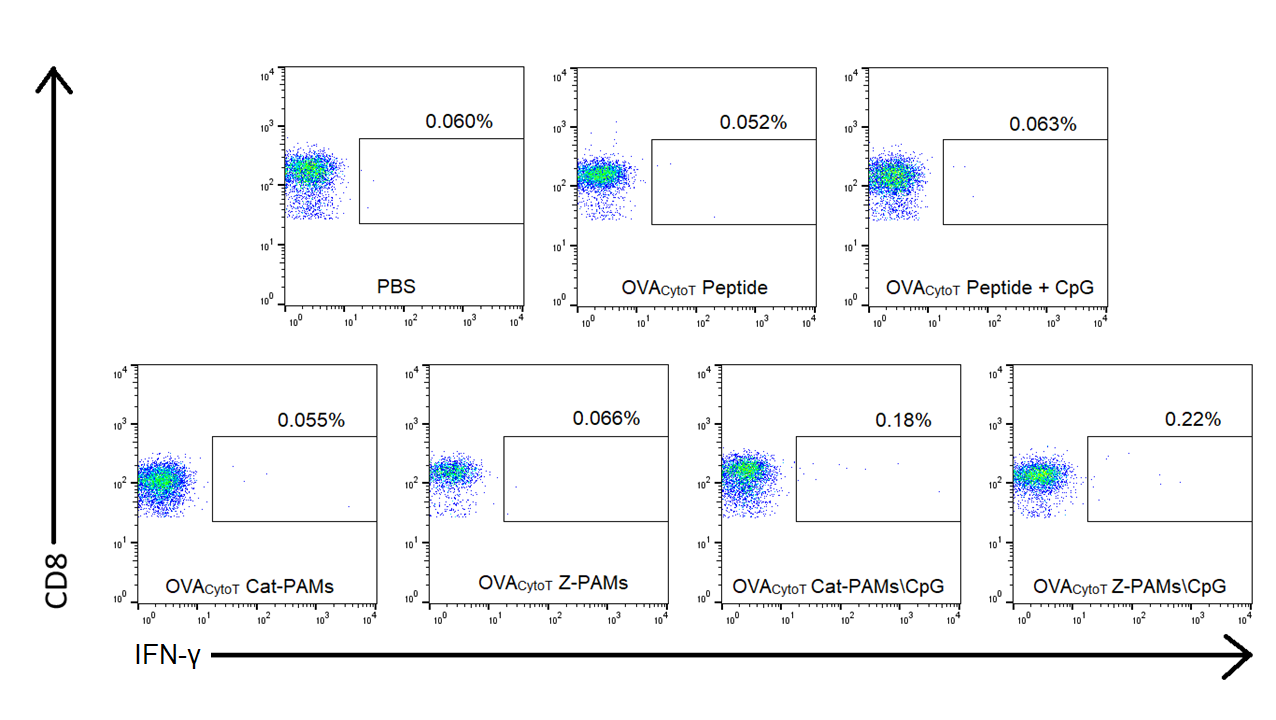

**Figure S3.** Representative lymph node CD8 versus IFN-γ flow cytometry dot plots. Population gating was compensated by utilizing no stain and single stain controls after which the data was analyzed by Flo Jo software. The presence of cell surface CD8 was first gated to separate out the cytotoxic T cell population from other living lymphocytes. This population was then investigated for the presence of cell associated IFN-γ to determine the percentage of IFN-γ^+^ CD8^+^ T cells.

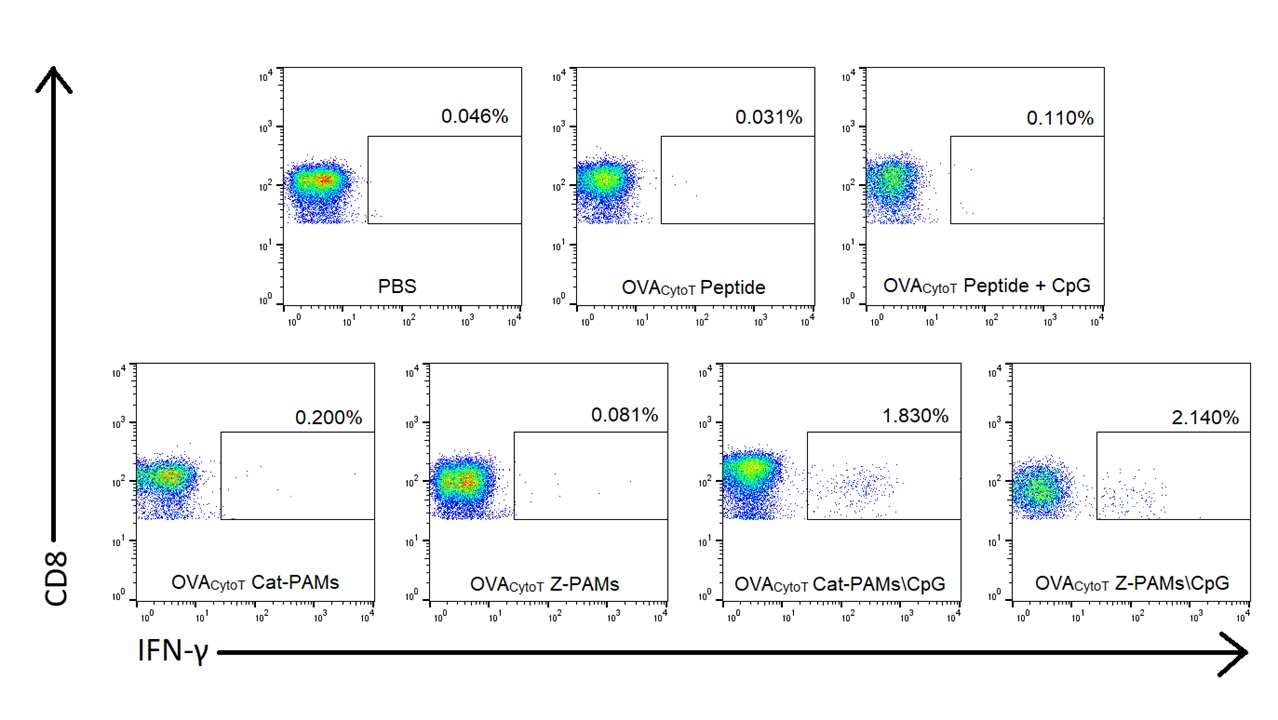

**Figure S4.** Representative spleen CD8 versus IFN-γ flow cytometry dot plots. Population gating was compensated by utilizing no stain and single stain controls after which the data was analyzed by Flo Jo software. The presence of cell surface CD8 was first gated to separate out the cytotoxic T cell population from other living lymphocytes. This population was then investigated for the presence of cell associated IFN-γ, thus determining the percentage of IFN-γ^+^ CD8^+^ T cells.

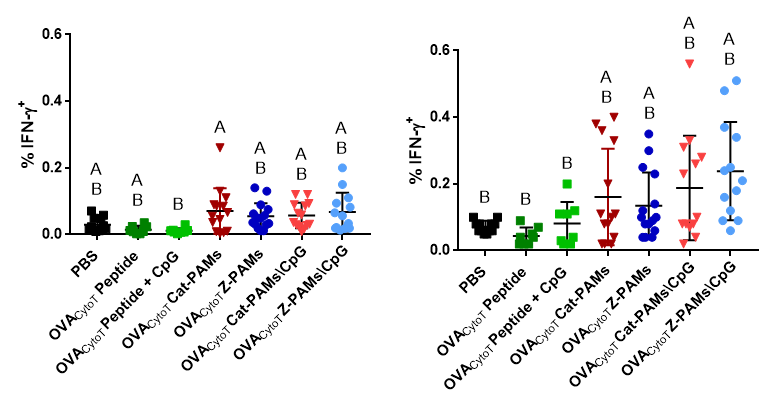

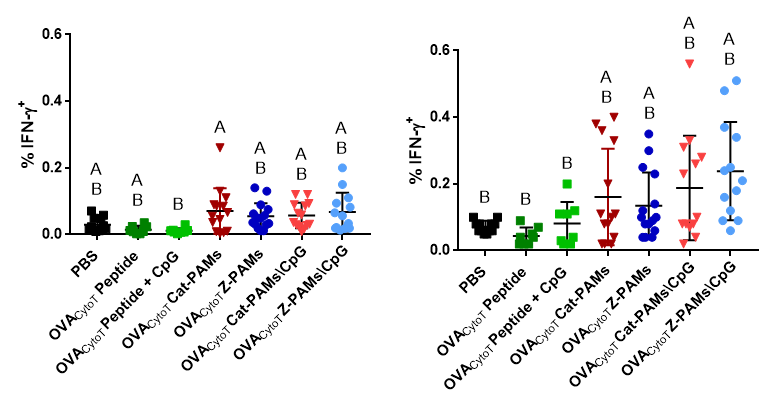

**Figure S5.** CpG complexed micelles did not enhance the immune response of CD8^-^ lymphocytes or splenocytes. Vaccine formulation didn’t affect antigen responsive of CD8^-^ cells in the **a** draining lymph nodes and **b** spleen. Specifically, co-delivery of OVA_CytoT­_ PAMs and CpG didn’t enhance the amount of IFN-γ^+^ CD8^-^ cells. Within the graph (For a and b), groups that possess different letters have statistically significant differences in mean (p ≤ 0.05) whereas those that have the same letter have similar means (p > 0.05).

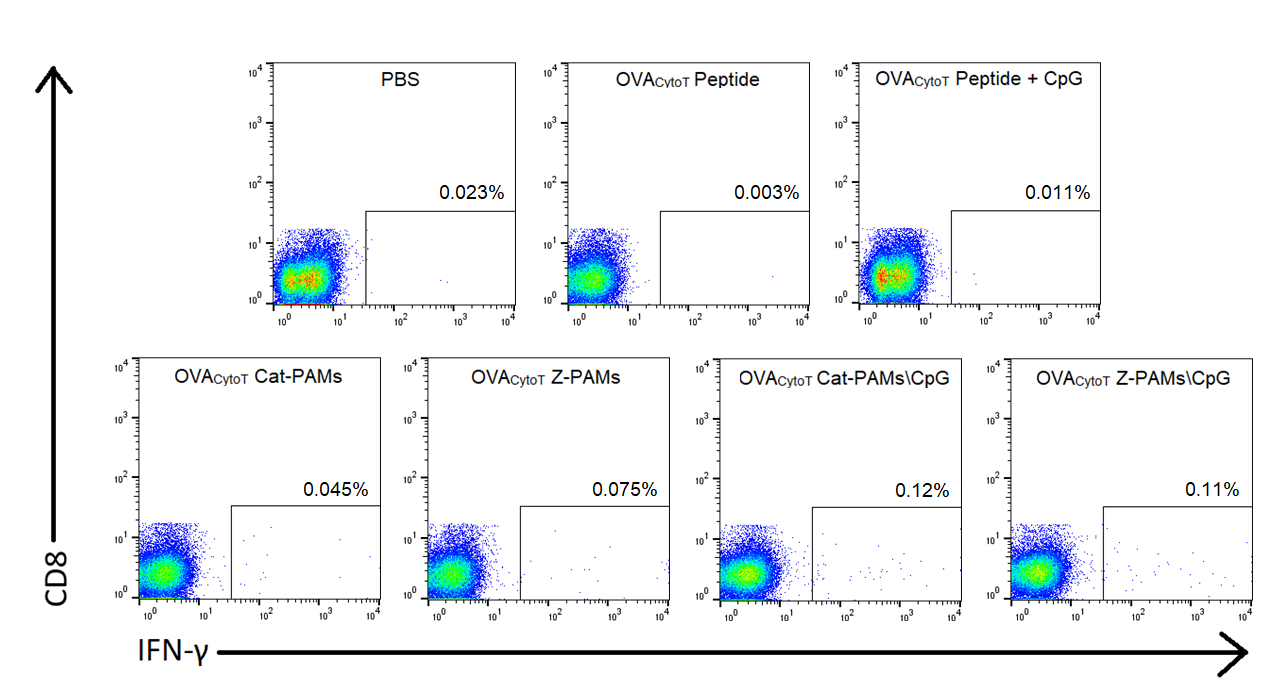

**Figure S6.** Representative lymph node CD8 versus IFN-γ flow cytometry dot plots. Population gating was compensated utilizing no stain and single stain controls after which the data was analyzed by Flo Jo software. The presence of cell surface CD8 was first gated to separate out the cytotoxic T cell population from other living lymphocytes. This population was then investigated for the presence of cell associated IFN-γ thus determining the percentage of IFN-γ^+^ CD8^-^ T cells.

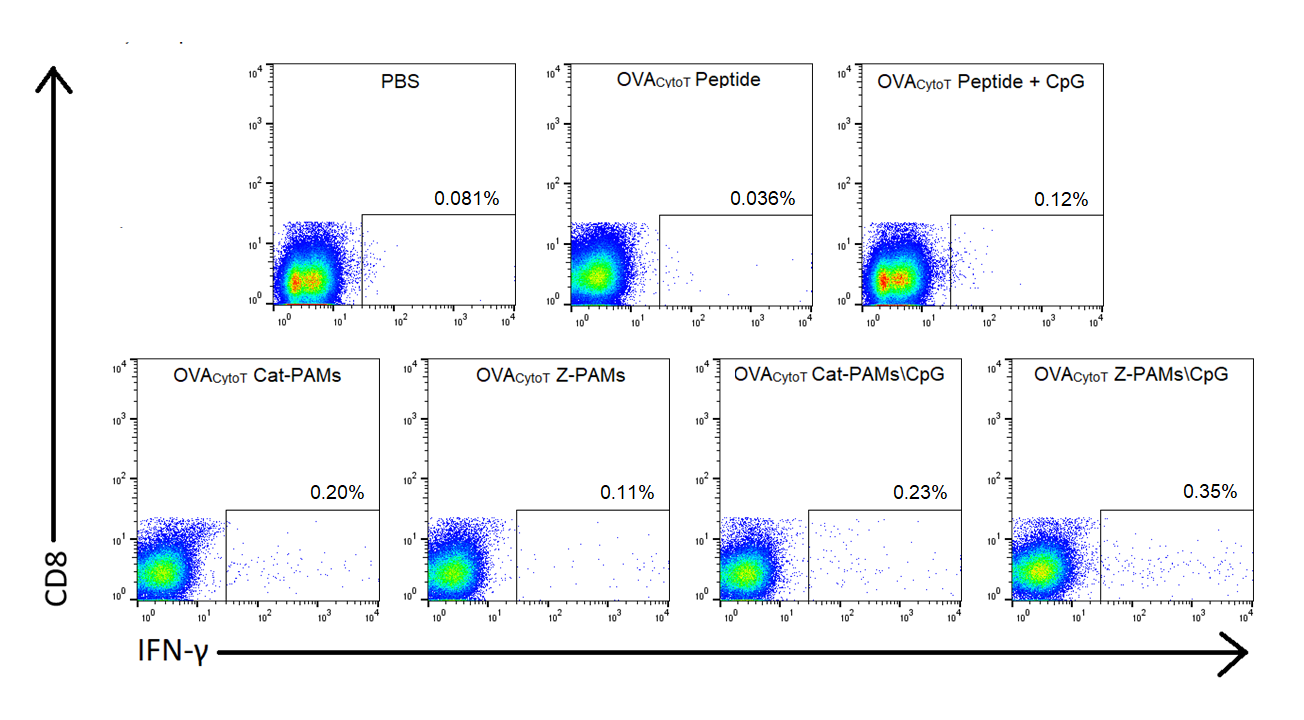

**Figure S7.** Representative spleen CD8 versus IFN-γ flow cytometry dot plots. Population gating was compensated utilizing no stain and single stain controls after which the data was analyzed by Flo Jo software. The presence of cell surface CD8 was first gated to separate out the cytotoxic T cell population from other living lymphocytes. This population was then investigated for the presence of cell associated IFN-γ thus determining the percentage of IFN-γ^+^ CD8^-^ T cells.

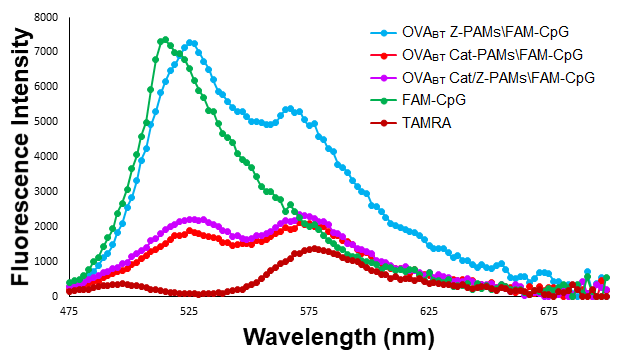

**Figure S8.** CpG readily complexes to charge matched and highly positively charged OVA_BT_ PAMs. Fluorescent spectra data from a Förster Resonance Energy Transfer (FRET) study show the presence of only the donor peak (~ 525 nm) for FAM-CpG and the acceptor peak (~ 580 nm) for TAMRA entrapped in OVA­_BT_ PAM formulation. When both OVA_BT_ Cat-PAMs and OVA_BT_ Cat/Z-PAMs were mixed with CpG during fabrication, the resulting formulations (i.e. OVA_BT_ Cat-PAMs\CpG and OVA_BT_ Cat/Z-PAMs\CpG) underwent electrostatic complexation as demonstrated by a considerable reduction and small enhancement in donor and acceptor fluorescence, respectively. The FRET study additionally provided evidence that OVA­_BT­_ Z-PAMs did not electrostatically complex with CpG as demonstrated by the similar fluorescent peak as the FAM-CpG control.
